# Co-inoculation with novel nodule-inhabiting bacteria reduces the benefits of legume-rhizobium symbiosis

**DOI:** 10.1101/2023.11.15.567292

**Authors:** James C. Kosmopoulos, Rebecca T. Batstone-Doyle, Katy D. Heath

## Abstract

The ecologically and economically vital symbiosis between nitrogen-fixing rhizobia and leguminous plants is often thought of as a bi-partite interaction, yet studies increasingly show the prevalence of non-rhizobial endophytes (NREs) that occupy nodules alongside rhizobia. Yet, what impact these NREs have on plant or rhizobium fitness remains unclear. Here, we investigated four NRE strains found to naturally co-occupy nodules of the legume *Medicago truncatula* alongside *Sinorhizobium meliloti* in native soils. Our objectives were to (1) examine the direct and indirect effects of NREs on *M. truncatula* and *S. meliloti* fitness, and (2), determine whether NREs can re-colonize root and nodule tissues upon reinoculation. We identified one NRE strain (522) as a novel *Paenibacillus* species, another strain (717A) as a novel *Bacillus* species, and the other two (702A and 733B) as novel *Pseudomonas* species. Additionally, we found that two NREs (Bacillus 717A and Pseudomonas 733B) reduced the fitness benefits obtained from symbiosis for both partners, while the other two (522, 702A) had little effect. Lastly, we found that NREs were able to co-infect host tissues alongside *S. meliloti*. This study demonstrates that variation of NREs present in natural populations must be considered to better understand legume-rhizobium dynamics in soil communities.

## INTRODUCTION

Plant-microbe symbioses have often been studied as pairwise interactions between a single plant species and a single species of bacteria or fungi (Bakker et al. 2014; Tsiknia et al. 2020; Afkhami et al. 2020). Yet, natural microbial communities are diverse in their functional roles, assembly processes, and evolutionary dynamics. Our understanding of this diversity has been growing thanks to a recent boom in plant microbiome studies (Kent and Triplett 2002; Wagner et al. 2014, 2016; Zilles et al. 2016; O’Brien et al. 2021). Plants in nature never interact with a single species at a time. Intimate symbiotic interactions (including plant-fungi or legume-rhizobium interactions) are occurring in a complex web of microbe-microbe (and other) interactions (Bakker et al. 2014). It has long been recognized that the impacts symbiotic partners have on one another are context-dependent, varying from beneficial (mutualistic), to commensal, to even harmful (parasitic) depending on abiotic and biotic contexts (Johnson et al. 1997; Heath and Tiffin 2007; Haney et al. 2015; Klein et al. 2022; Batstone et al. 2022b). Studying symbiotic outcomes in light of the broader microbiome community will ultimately lead to a more predictive understanding for how intimate plant-microbe symbioses evolve in real ecosystems (Tsiknia et al., 2020).

The model symbiotic mutualism between leguminous plants and the nitrogen fixing rhizobial bacteria (rhizobia) is responsible for significant contributions to bioavailable N sources in unmanaged and agricultural systems (Vitousek et al. 1997; Herridge et al. 2008b). In this interaction, legumes house rhizobia within nodules that develop on their roots and provide rhizobia with photosynthate, while in return rhizobia reduce atmospheric nitrogen N_2_ to ammonia NH_3_ (Gage, 2004). With the advent of next generation sequencing, it has become increasingly clear that rhizobia are not the only endophytic occupants of root nodules. Nodules can be colonized by non-rhizobium endophytes (NREs), bacteria that come from dozens of genera and span multiple phyla (Martínez-Hidalgo et al. 2014; Martínez-Hidalgo and Hirsch 2017; Tokgöz et al. 2020; Rahal and Chekireb 2021). Of the NREs that have been described so far, co-inoculation has resulted in either positive or neutral effects on plant growth and nodule phenotypes (Khan, 2019; Martínez-Hidalgo & Hirsch, 2017). For example, *M. truncatula* co-inoculated with *Sinorhizobium medicae* WSM419 and *Pseudomonas fluorescens* WSM3457 formed a greater number of nodules (Fox et al. 2011). Also, *Medicago sativa* co-inoculated with *Sinorhizobium meliloti* 1021 and *Micromonospora* isolates had more efficient nutrient uptake and increased shoot mass (Martínez-Hidalgo et al., 2014). Thus, based on the limited number of studies to date, NREs have often been broadly described as plant-growth promoting bacteria (PGPBs) (Martínez-Hidalgo & Hirsch, 2017). However, the NREs tested thus far were chosen because of their plant-growth promoting potential on other non-legumes, meaning they are likely to be biased towards showing beneficial impacts. It remains unclear whether the diversity of NREs that naturally co-occur with rhizobia and plant genotypes will similarly have positive effects on symbiotic outcomes, or whether some might have negative impacts.

Shared coevolutionary history and/or shared environmental conditions between partners can heavily influence the effects of symbiosis on hosts. For example, populations of *Mesorhizobium* associated with *Acmispon wrangelianus* legume hosts displayed contrasting levels of heavy metal adaptation depending on shared soil type (Porter et al. 2017). Additionally, plant-associated microbial communities became more beneficial for their plant hosts under the appropriate drought regime (Lau and Lennon 2012; Bolin et al. 2023). In one study, legume genotypes inoculated with rhizobia that had been experimentally evolved on the same genotype grew better compared to those inoculated with rhizobia that evolved on different host genotypes (Batstone et al., 2020). Some NREs could have positive impacts on hosts and rhizobia if they increase the net benefits gained by partners, for example by providing rhizobia with a critical metabolite that enhances their ability to fix N. Other NREs might instead act as rhizobium competitors and/or plant pathogens and thereby reduce the benefits exchanged and thus the fitness of host and/or rhizobia. Predicting the ecologically relevant impacts that NREs have on legume and rhizobia in symbiosis will require a direct comparison of the net benefits gained by both partners in the presence or absence of multiple, distinct NREs that co-occupy nodules collected from hosts in native soils.

Here, we use four naturally occurring NREs and a strain of *Sinorhizobium meliloti* (strain 141; Batstone et al. 2022b) (formerly *Ensifer meliloti*) that were isolated from soils in the native range of the model legume *Medicago truncatula* (Riley et al. 2023). Our objectives were to (1) compare fitness proxies of both plant (e.g., above-ground shoot biomass) and rhizobium (e.g., nodule number) when plants were inoculated with *S. meliloti* 141 alone versus plants co-inoculated with both *S. meliloti* and one of the four NREs; and (2), verify whether NREs could be found in root and nodule tissues of plants when co-inoculated with *S. meliloti* 141.

## METHODS

### Study system

We chose the *Medicago truncatula* genotype DZA 315.6 (hereafter DZA) because we previously characterized its growth when paired with hundreds of rhizobial strains (Batstone et al. 2022a, 2022b), and it has a well-documented ability to form root nodules with diverse soil endophytes (Etienne-Pascal & Pascal, 2013). We chose *Sinorhizobium meliloti* strain 141 (hereafter *Sinorhizobium*) as our focal rhizobium partner because it was found to be a high-quality symbiont (conferring a high fitness benefit to its host) across several *Medicago truncatula* genotypes (A17 and DZA; Batstone et al. 2022b). The four non-rhizobial endophytes (NRE) used in this study, along with *Sinorhizobium*, were previously isolated from the root nodules of natural populations of *Medicago truncatula*, as described in detail in Riley et al. (2023). Briefly, strains were isolated from soils surrounding *M. truncatula* roots from 21 sites spanning the species’ native range: Spain, France, and Corsica. For more detail on how microbial strains were isolated and taxonomically characterized, see Supplemental Methods.Click or tap here to enter text.

### Whole-genome species trees

We first assigned our four NREs to putative genera by aligning the 16S rRNA gene sequences of each against the NCBI RefSeq 16S rRNA database (O’Leary et al., 2016), and inferring genera based on the top five BLAST hits. Since NRE strain 522 contained 16S sequences of both *Paenibacillus* and *Brevibacillus* origin (see Results), *Brevibacillus* contigs in the genome assembly for strain 522 were removed using VizBin (Laczny et al. 2015) before further analysis. Because species-level identification of bacteria often requires a high-resolution phylogenetic analysis (Lan et al., 2016), we constructed minimum-evolution phylogenomic trees based on whole-genome alignments using the Type (Strain) Genome Server (Meier-Kolthoff and Göker 2019) (Supplemental Methods). For the phylogenetic inference, we conducted all pairwise comparisons among the set of genomes using the Genome BLAST Distance Phylogeny approach (GBDP) (Meier-Kolthoff et al. 2013) and we inferred accurate intergenomic distances under the algorithm ‘trimming’ and distance formula *d_5_*. One-hundred distance replicates were calculated each. We calculated digital DDH values and confidence intervals using the recommended settings of the GGDC 4.0 (Meier-Kolthoff et al. 2013, 2022). We used the resulting intergenomic distances to infer a balanced minimum evolution tree with branch support via FASTME 2.1.6.1 including SPR postprocessing (Lefort et al. 2015). We inferred branch support from 100 pseudo-bootstrap replicates each, and we rooted trees at the midpoint. We performed type-based species clustering using a 70% dDDH radius around each of the 17 type strains as previously described (Meier-Kolthoff and Göker 2019). The resulting groups are shown in Table S1 and Table S2. We clustered subspecies using a 79% dDDH threshold as previously introduced (Meier-Kolthoff et al. 2014).

### Plant genotype and growth methods

We razor scarified and surface-sterilized seeds of *M. truncatula* genotype DZA, washing them in 95% ethanol for 30 seconds and then commercial bleach for seven minutes, followed by sterile water for four minutes to rinse off any excess bleach. Before planting, we packed the bottoms of SC10R Ray Leech “Cone-tainers” (Stuewe & Sons, Inc., Tangent, OR) with a small handful of autoclaved polyester fiber to cover their drainage holes. We filled each Cone-tainer with ∼ 200 mL of an autoclave-sterilized mixture of one-part root wash: one-part sand: four-parts turface MVP calcined clay (Profile Products LLC, Buffalo Grove, IL, USA). The resulting potting media composition is suitable for *M. truncatula* growth because it facilitates root extraction and rinsing upon harvest, which greatly expediates measuring nodule phenotypes (see below). We sowed seeds ¾ centimeter deep in the potting media-filled Cone-tainers using sterile forceps, and immediately watered to compact the soil and prevent seed desiccation. We sowed two seeds in each pot in case one failed to germinate, and then covered the potting media surface and seeds with ½ cm of autoclaved-sterilized vermiculite to help retain moisture around seeds after sowing. Prior to inoculation, we thinned all pots to one seedling using sterile forceps.

### Inocula preparation

For each of our five strains (four NREs and one *Sinorhizobium*), we separately streaked out stocks onto sterile petri dishes with solid Tryptone-Yeast (TY) medium (Vincent, 1970). We allowed colonies to grow in a 30 °C dark incubator until ample growth was observed, approximately 48 hours. For each strain, we picked a single colony to inoculate a sterile 15 mL Falcon tube filled with liquid TY medium, and then placed the capped tubes in a shaking incubator set to 30 °C and 200 rpm for overnight growth. Between 20-24 hours later, we combined tubes of the same strain into a single sterile 50 mL falcon tube, gently inverted to mix, and pipetted out 500 μL to estimate cell density (cells per mL) via measuring absorbance at OD_600_ using a NanoDrop 2000c (Thermo Scientific; Waltham, MA, USA). To ensure that each strain started at an equal inoculation density, we added an appropriate amount of sterile liquid medium and culture to reach a final OD_600_ of 0.1, which corresponds to ∼1 x 10^8^ cells/mL.

### Greenhouse experiment

We tested the direct and indirect effects of NREs on the legume-rhizobium symbiosis by comparing co-inoculations of *Sinorhizobium* with each of the four NREs to single-inoculations with *Sinorhizobium* alone or each of the four NREs alone, for a total of 10 treatments: five single-inoculation, four co-inoculation, plus one uninoculated control to monitor contamination levels. The single-inoculation treatments consisted of either a 500 μL dose of *Sinorhizobium* (hereafter “*Sinorhizobium*-only”) or a 500 μL dose of one of the four NREs (hereafter “NRE-only”), each dose totaling ∼5x10^7^ cells (1x10^8^ cells/mL x 0.5 mL). The co-inoculation treatments consisted of one full 500 μL dose of one of the four NRE strains *and* a full 500 μL dose of *S. meliloti* 141 (hereafter “co-inoculation”), the combined dose totaling 1x10^8^ cells (1x10^8^ cells/mL x 1 mL). Given that *Sinorhizobium* was the plant’s only source of fixed N and was thus expected to be a major limiter on plant growth regardless of NRE presence, we opted to control for the number of *Sinorhizobium* cells across inoculation treatments rather than the total number of cells. The positions of the racks that held *Medicago* plants were randomized within the greenhouse (Supplemental Methods). We inoculated plants with their respective treatments seven days after seeds had been sown (i.e., when the first true leaf had emerged) to give the plants sufficient time to establish their root systems and begin photosynthesizing. Plants were destructively harvested four weeks post inoculation (Supplemental Methods).

### Nitrogen addition experiment

For NREs that were found to significantly impact plant traits, we wanted to tease apart whether these effects were acting on the plants directly (i.e., observed in the absence of co-inoculation with rhizobia) or indirectly (i.e., via their interactions with rhizobia). However, because we did not supply plants with any external sources of N in our greenhouse experiment (described above), plants grew poorly in all treatments in which *Sinorhizobium* was absent, precluding our ability to thoroughly test direct vs indirect effects of NREs on plant growth. To address this limitation, we conducted an additional experiment in which plants were supplied with moderate amounts of N and were either inoculated with a single NRE or were left uninoculated (control). If plants grew larger or smaller when inoculated with an NRE compared to uninoculated controls, then we would consider this NRE to incur direct benefits (i.e., mutualistic) or costs (i.e., pathogenic) to the plant, respectively. We prepared 16 replicate plants per inoculation treatment, using similar protocols as described for the greenhouse experiment (above). Briefly, plants were grown in magenta boxes (PlantMedia, Dublin, OH) and were placed in a growth chamber and were treated with N-supplemented media once a week. One week after transplanting, plants were either inoculated with each NRE individually or with sterile media. After four weeks, we destructively harvested plants as described above. For more detail on the nitrogen addition experiment, see Supplementary Methods.

### Statistical analyses on phenotypic data

For data measured in the greenhouse experiment, we constructed a linear mixed-effects model (LMM) using the package “lme4” (Bates et al., 2015) in R (R Core Team, 2020) to first test whether measured traits were influenced by inoculation treatments. For each model, we included the fixed effects of inoculation treatment (10 levels) and whether plants were inoculated with rhizobia or not (2 levels: present/absent) to account for effects varying between the two groups. Plant rack was included as a random effect to account for the spatial arrangement of racks in the greenhouse. We performed type II ANOVA on the model generated for each trait using the “lmerTest” package (Kuznetsova et al. 2017). We were most interested in testing how traits responded to co-inoculations with NREs versus those without; thus we used the “emmeans” package (Lenth 2022) to estimate the marginal means of each treatment group for plants inoculated or uninoculated with rhizobia separately. To compare among groups, we used the Dunnet-adjusted “trt.vs.ctrl” option to compare the treatment groups to two different controls: for plants uninoculated with rhizobia, treatment groups were considered to be plants inoculated with each of the four NREs individually while the control was uninoculated plants. For plants inoculated with rhizobia, treatment groups were co-inoculated plants (rhizobia + each NRE) while the control was plants inoculated with rhizobia only. Click or tap here to enter text.For the nitrogen experiment, we analyzed data using LMMs and a similar post-hoc analysis as described above, but with inoculation treatment (control, inoculated) as fixed effect and pot position as a random effect. For each model, we performed residual diagnostics to check for heteroskedasticity and non-independence, and decided that transforming our response variables was not required.

### Tissue occupancy experiment

Finally, we wanted to confirm whether NREs that significantly impacted the symbiosis, which were originally isolated from nodules that formed on plants growing in the field, could reinfect plant tissues after inoculation. We conducted an additional co-inoculation experiment of *Sinorhizobium* paired with each NRE that affected host growth (see Results). Seeds of *M. truncatula* DZA were prepared, inoculated, and harvested using the same methods as the nitrogen addition experiment (see above), and with slight deviations (Supplemental Methods). Following the harvest, we removed nodules from roots as well as root tissue sections without nodules. We then partitioned these tissue samples for DNA sequencing into two categories: endophytes (within tissues) as well as endophytes + epiphytes (on tissue surfaces). In total, we gathered 48 tissue samples for DNA sequencing (3 treatments x 2 tissue sections x 2 sterilization treatments x 4 replicates = 48 samples). To sequence only endophytic bacteria, we randomly chose four samples from each tissue section per inoculum treatment and surface sterilized each individually by adding 1 mL of 30% commercial bleach, thoroughly mixing for 60 seconds, and then removing bleach thoroughly by washing each sample with 1 mL of sterile DI water. To sequence epiphytic in addition to endophytic bacteria, we randomly chose four additional samples per tissue section per treatment, washing each individually with 1 mL of sterile DI water, and thoroughly mixing for 60 seconds to remove loosely-associated bacteria on tissue surfaces.

### DNA extraction and 16S V3-V4 amplicon sequencing

We extracted and purified genomic DNA from tissue samples using a DNeasy Plant Pro Kit (Qiagen, Hilden, Germany) following the manufacturer’s instructions. For each sample, we evaluated the DNA concentration using optical density measurements obtained by a Nanodrop spectrophotometer (NanoDrop Technologies, Rockland, DE, USA) at 260 and 280 nm wavelengths. We submitted DNA samples to the W.M. Keck Center for Comparative and Functional Genomics at the University of Illinois at Urbana-Champaign for 16S rRNA V3-V4 region amplicon sequencing. Briefly, DNA quality was assessed with a Qubit fluorometer (Invitrogen, MA, USA) and the targeted 16S regions were amplified by PCR using primers V3-F357_N (5’-CCTACGGGNGGCWGCAG-3′) and V4-R805 (5’-GACTACHVGGGTATCTAATCC-3′) using a Fluidigm Biomark HD PCR machine (Fluidigm Corporation, South San Francisco, CA, USA). PCR products were quantified with a QuantiT PicoGreen fluorometer (Invitrogen, MA, USA), then cleaned, purified, and size-selected in a 2% agarose E-gel (Invitrogen, MA, USA) followed by gel extraction with a QIAquick Gel extraction kit (Qiagen, Hilden, Germany). Amplicons were sequenced with Illumina MiSeq v2 platform (Illumina, San Diego, CA, USA) according to the manufacturer’s instructions.

### Amplicon sequence variant (ASV) inference

For each library (i.e., sample), we inspected raw, demultiplexed reads for quality using DADA2 v1.28.0 (Callahan et al. 2016) in R (R Core Team 2020). Reads were trimmed with DADA2 to remove forward and reverse primers at lengths of 32 and 35, respectively, to account for the length of V3-V4 primers plus the length of Fluidigm-specific CS primers. We filtered the resulting trimmed reads with DADA2 to retain reads with a maximum number of expected errors of two on the forward strand and five on the reverse strand, in addition to removing putative phiX sequences. We truncated reads at the first instance of a quality score less than or equal to two. The resulting filtered and trimmed reads were dereplicated with DADA2 and amplicon sequence variants (ASVs) were inferred from the dereplicated reads with DADA2. We merged ASVs from forward and reverse read pairs and removed putative chimeras with DADA2, using the “consensus” method. The resulting merged, non-chimeric ASVs were used to construct a sequence table, and the taxonomy of the ASVs was inferred with DECIPHER v2.28.0 (Wright 2016) with the SILVA SSU r138 database (Quast et al. 2012). We manually renamed the resulting taxonomic assignments to the genus *Ensifer* to *Sinorhizobium* to reflect the most recent modifications to the genera *Ensifer* and *Sinorhizobium* (Kuzmanović et al. 2022) Additionally, we removed ASVs without a taxonomic assignment at the domain level and ASVs assigned to the orders Rickettsiales or “Chloroplast” using phyloseq v1.44.0 (McMurdie and Holmes 2013) to exclude ASVs inferred from host DNA from downstream analyses. We constructed a heatmap visualizing the matrix of variance-stabilized ASVs counts per sample using the R package pheatmap v1.0.12 (Kolde 2019) using color palettes generated by RColorBrewer v1.1.3 (Neuwirth 2022) and viridis v0.6.3 (Garnier et al. 2023). The resulting heatmap was manually edited to position color legends and to format the font face of ASV labels for Figure 4. ASV names with polyphyletic or *Candidatus* genus assignments from SILVA were manually replaced with “NA”.

### ASV differential abundance analysis

To determine the relative enrichment of ASVs across inoculum treatment, tissue section, and surface-sterilization treatment, we used the absolute, non-normalized ASVs counts in conjunction with DESeq2 (Love et al. 2014) v1.40.2 for a differential abundance analysis. Briefly, we built negative binomial generalized models using the ASV counts in DESeq2 with inoculum, tissue section, and surface-sterilization treatment included as factors in the models. To determine ASVs whose abundances were significantly affected by each factor individually, we performed likelihood ratio tests using DESeq2 using full models of the three factors together and reduced models lacking one of each factor. For each test, we shrunk the resulting log2 fold-changes of every contrast of the factor’s levels using the DESeq2 function “lfcShrink”. Lastly, to determine ASVs differentially abundant across surface-sterilization treatments among the same tissue sections, we performed an additional likelihood ratio test. The full model in this test contained one factor representing the interaction of tissue section and surface-sterilization treatment, while the reduced model included only the intercept. As done with the other tests, we shrunk the log2 fold changes of each level contrast. We generated all plots with ggplot2 (Wickham 2016).

### ASV BLAST alignments

Using NCBI BLAST+ v2.14.0, we created a nucleotide database from the 16S rRNA sequences annotated from the genome assemblies of *Sinorhizobium*, *Bacillus* sp. 717A, and *Pseudomonas* sp. 733B. All ASV nucleotide sequences inferred by DADA2 (Table S7) were used as a query in a BLASTn search with BLAST+ (Camacho et al. 2009), retaining alignments with a minimum e-value of 0.01 and at least 80% identity. ASV28 (Table S7) was determined to be a likely representative of *Bacillus* sp. 717A (see results) and was aligned against the NCBI rRNA type strain database on the BLAST+ online platform (blast.ncbi.nlm.nih.gov) to verify its potential taxonomic assignment. We determined that ASV1 and ASV5 were representatives of *Sinorhizobium* 141 and *Pseudomonas* sp. 733B, respectively (see results). To investigate the presence of strains related to these two ASVs plus ASV28 in other contexts, we aligned ASV1, ASV5, and ASV28 against the 50 most abundant sequences inferred from 16S rRNA amplicon sequencing in another study of symbiotic bacteria of *M. truncatula* (Brown et al. 2020) using BLASTn with the same parameters.

## RESULTS

### Taxonomy

Our 16S rRNA BLAST results suggested NRE strain 522 belongs to the genus *Paenibacillus*, 717A belongs to *Bacillus*, while 702A and 733B belong to *Pseudomonas*. This process revealed that the whole-genome sequence for *Paenibacillus sp.* 522 was contaminated with contigs from a *Brevibacillus*, and so *Brevibacillus* contigs were removed from the 717A genome assembly before tree construction to avoid errors; however, it is possible that this *Brevibacillus* was present in the 522 inoculations. We subjected all strains to minimum-evolution species tree construction using whole-genome sequences. Based on the inferred phylogenomic trees (Figure 1) and the calculated genome distances between the query and type strains (Tables S1-S2), we identified *Paenibacillus* sp. 522 (Figure 1A), *Bacillus* sp. 717A (Figure 1B), *Pseudomonas* spp. 702A and 733B (Figure 1C) each as novel species in their genera.

**Figure 1.**
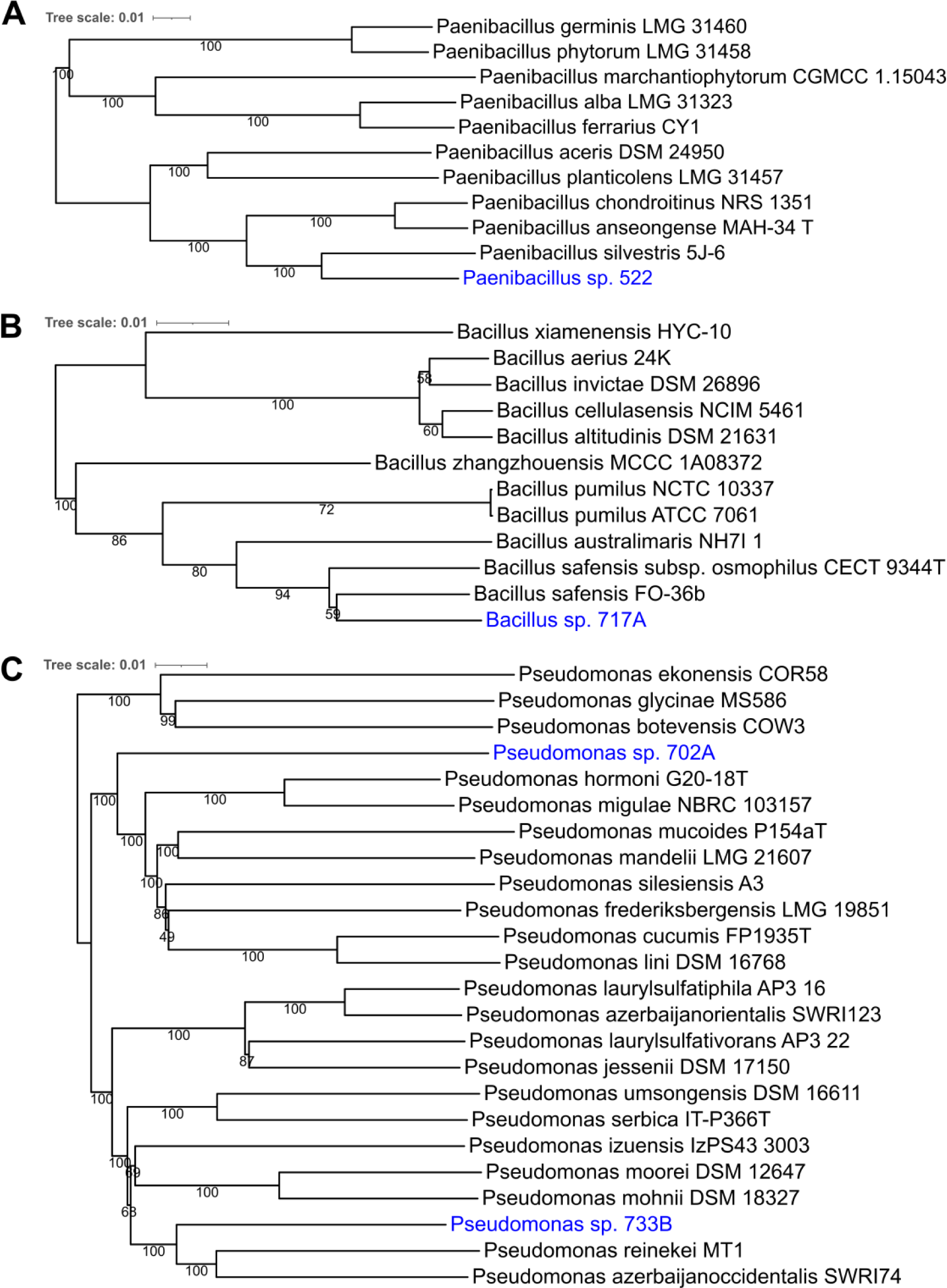
Novel *Paenibacillus, Bacillus,* and *Pseudomonas* species among non-rhizobia endophytes. Whole-genome phylogenies inferred by the Type (Strain) Genome Server (Meier-Kolthoff and Göker 2019). Trees inferred with FastME 2.1.6.1 (Lefort et al. 2015) from GBDP distances calculated from genome sequences of strains 522 (A), 717A (B), 702A and 733B (C). The branch lengths are scaled in terms of the Genome BLAST Distance Phylogeny (GBDP) formula *d_5_* (Meier-Kolthoff et al. 2013). The numbers at branches are GBDP pseudo-bootstrap support values > 60% from 100 replications, with an average branch support of and 100% (A), 78.8% (B), and 93.2% (C). All trees were rooted at the midpoint. Blue taxa are query non-rhizobial endophyte strains, black taxa are type strains from the TYGS database (Meier-Kolthoff et al. 2022).

### Greenhouse experiment

We found no clear impact on shoot mass between plants inoculated with *Sinorhizobium* alone compared to plants co-inoculated with *Sinorhizobium* and either *Paenibacillus sp.* 522 or *Pseudomonas sp.* 702A, whereas plants coinoculated with *Sinorhizobium* plus either *Bacillus sp.* 717A or *Pseudomonas sp.* 733B grew significantly more poorly compared to *Sinorhizobium*-only plants (*P*-values < 0.05), with a 37% and 33% reduction in shoot mass, respectively (Figure 2A; Table S4). Every group of plants singly inoculated with an NRE without N-fixing *Sinorhizobium* and uninoculated control plants were in observably poor condition upon harvest (shoot mass ranging from ∼36-46 mg on average compared to ∼166-263 mg for plants inoculated with rhizobia; Table S5), with no clear difference in shoot mass between the NRE-only and uninoculated control groups (Figure 2A; Table S4).

**Figure 2.**
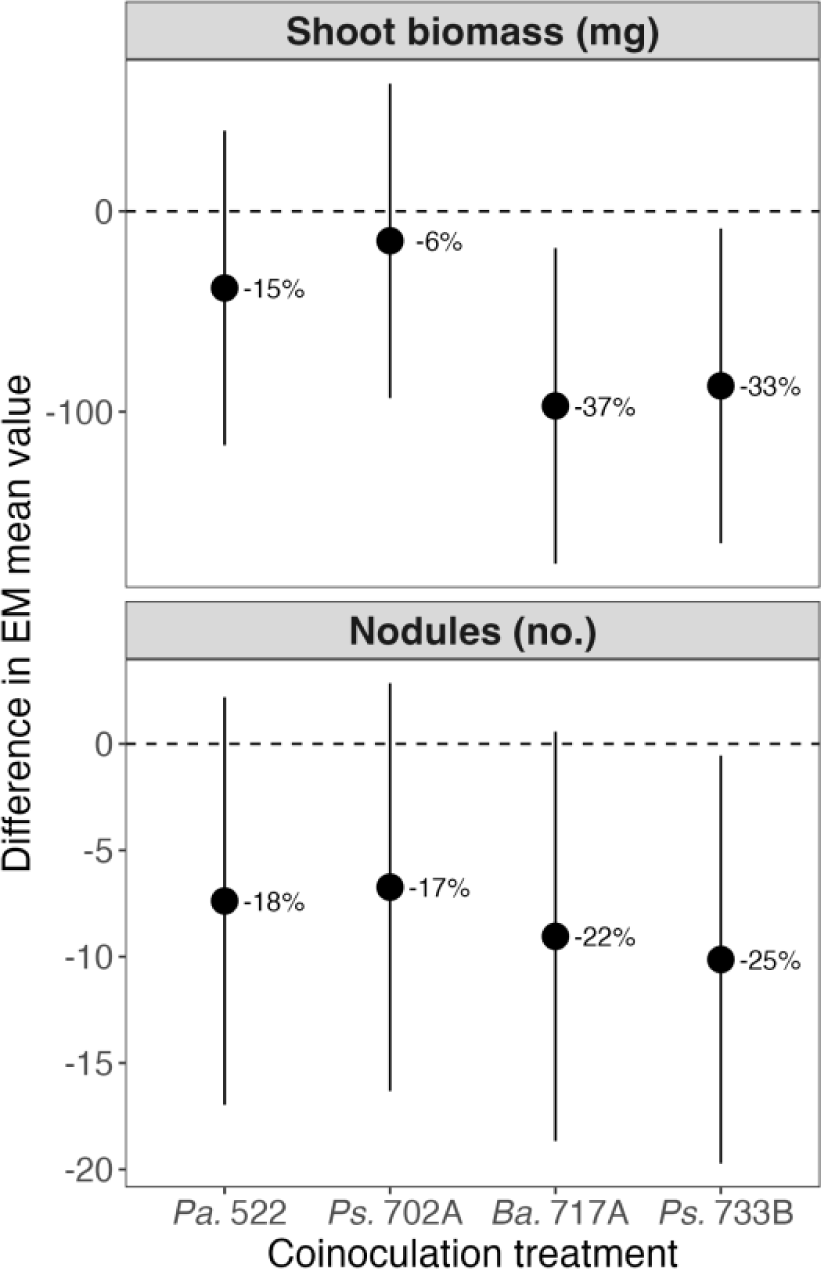
Co-inoculations of *Sinorhizobium meliloti* and *Bacillus* sp. 717A or *Pseudomonas sp.* 733B indirectly reduced host shoot mass and the number of nodules formed. Co-inoculations with *S. meliloti* and *Paenibacillus* sp. 522 or *Pseudomonas* sp. 702A did not yield different shoot masses (A) or numbers of nodules (B) than plants inoculated with *S. meliloti* only (*P* > 0.05, type II ANOVA, Dunnet adjusted), while co-inoculations with *Bacillus* sp. 717A or *Pseudomonas* sp. 733B significantly reduced the average shoot mass (A) and number of nodules (B) on plants (*P* < 0.01, *P* < 0.05, type II ANOVA, Dunnet adjusted). *Pa.* 522: *S. meliloti* 141 and *Paenibacillus* sp. 522 co-inoculation; *Ps.* 702A: *S. meliloti* 141 and *Pseudomonas* sp. 702A co-inoculation; *Ba.* 717A: *S. meliloti* 141 and *Bacillus* sp. 717A co-inoculation; *Ps.* 733B: *S. meliloti* 141 and *Pseudomonas* sp. 733B co-inoculation.

Uninoculated plants showed negligible signs of contamination; we only found two nodules on a single control plant, while all other controls (n = 29) were nodule-free. Similarly, we only found two plants inoculated with an NRE but not *Sinorhizobium* that formed nodules (one with 46 and the other with 31 nodules), which were subsequently removed from downstream analyses.

The lack of nodules on most plants (n = 78/80) that had been inoculated only with NREs (Figure 2B) indicated that these NREs were unable to form nodules on their own. Similarly to shoot mass, we found no clear impact on the average number of nodules formed when plants had been inoculated with *Sinorhizobium*-only versus those co-inoculated with *Paenibacillus sp.* 522 or *Pseudomonas sp.* 702A (Figure 2B; Table S4). However, both *Bacillus sp.* 717A and *Pseudomonas sp.* 733B co-inoculated plants formed significantly fewer nodules on average compared to *Sinorhizobium*-only plants, with a 22% and 25% reduction (n = 20), respectively, while none of the co-inoculated groups were significantly different from each other (Figure 2B; Table S4). This pattern is consistent with the differences observed in shoot mass, above. For a comprehensive list of all pairwise contrasts for each trait measurement and their associated *P*-values, see Table S4.

### Nitrogen addition experiment

While co-inoculations of *Sinorhizobium* with *Bacillus sp.* 717A or *Pseudomonas sp.* 733B negatively impacted host-symbiont traits (Figure 2B; Table S4), the effects on *M. truncatula* shoot mass could have been either an indirect result of an antagonistic interaction between NRE and *Sinorhizobium* or a direct result of parasitism by the NREs on the *M. truncatula* host. Since plants in the greenhouse experiment did not receive supplemental N, we could not distinguish whether the poor performance of *M. truncatula* plants singly inoculated with NREs (without N-fixing *Sinorhizobium*) was due to N-deficiency alone or if NRE inoculations also contributed. To address this, we tested whether the NREs were plant pathogens by examining the direct effects of *Bacillus sp.* 717A and *Pseudomonas sp.* 733B on host plants that received supplemental N fertilizer (Figure 3). There was no clear effect on growth when plants were inoculated with either strain compared to the uninoculated controls (*P* > 0.1, type II ANOVA), suggesting that the NREs 717A and 733B indirectly affected plant fitness in co-inoculations by inhibiting the legume-rhizobium symbiosis. Again plants grown without supplemental fertilizer (and without *Sinorhizobium*) were in poor condition (Table S6), and no nodules were observed on any plant.

**Figure 3.**
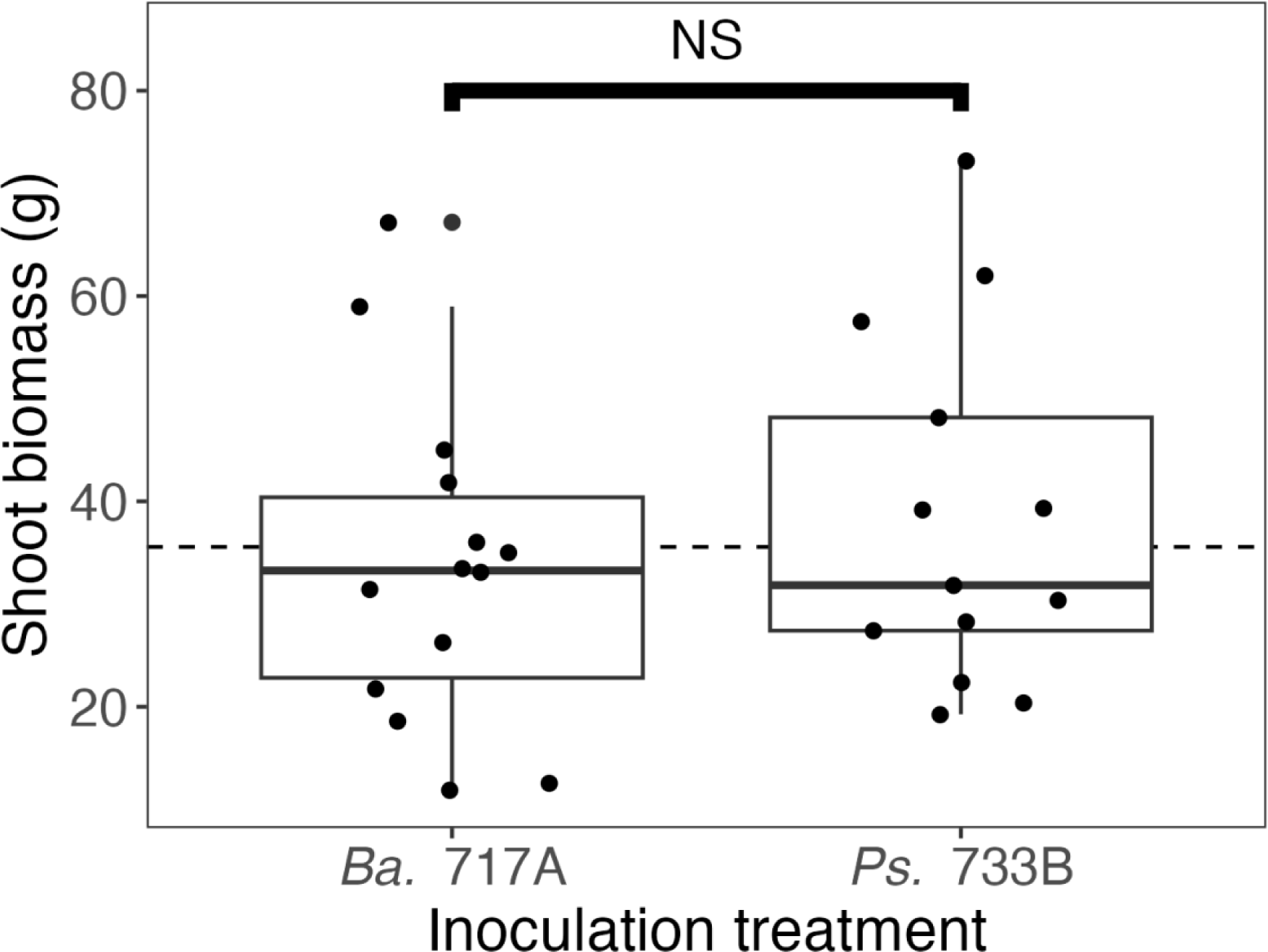
No evidence that NREs *Bacillus* sp. 717A and *Pseudomonas* sp. 733B had any direct effects on nitrogen-supplemented plants. Average shoot biomass for N-supplemented (NH_4_NO_3_) plants in the direct effects of NREs experiment are given. Inoculation treatment did not have a significant effect on shoot mass (*P* > 0.1, type II ANOVA). *Ba.* 717A: *Bacillus* sp. 717A single inoculation; *Ps.* 733B *Pseudomonas* sp. 733B single inoculation; NS: not significant.

### Tissue occupancy experiment

Finally, we used 16S rRNA amplicon sequencing of both surface-sterilized and non-surface-sterilized root and nodule tissues from coinoculated plants to locate these NREs in the endosphere and/or rhizosphere. Although we inoculated plants with combinations of only *Sinorhizobium*, *Bacillus sp.* 717A, and *Pseudomonas* sp. 733B in initially axenic conditions, taxonomic assignments of ASVs after four weeks in the greenhouse showed that other bacteria were also associated with our tissue samples (Figure 4). A total of 248 unique ASVs were inferred across all samples (Table S7); however, the vast majority of these ASVs were in very low abundance (Table S7). This low abundance is expected to some degree considering that 16S rRNA amplicon sequencing can report ASVs with abundances < 0.1% (Nikodemova et al. 2023), which either reflect spurious inferences or trace amounts of microbial cells. Nonetheless, ASVs were inferred from all inoculation, tissue, and surface-sterilization groups (Figure 4).

**Figure 4.**
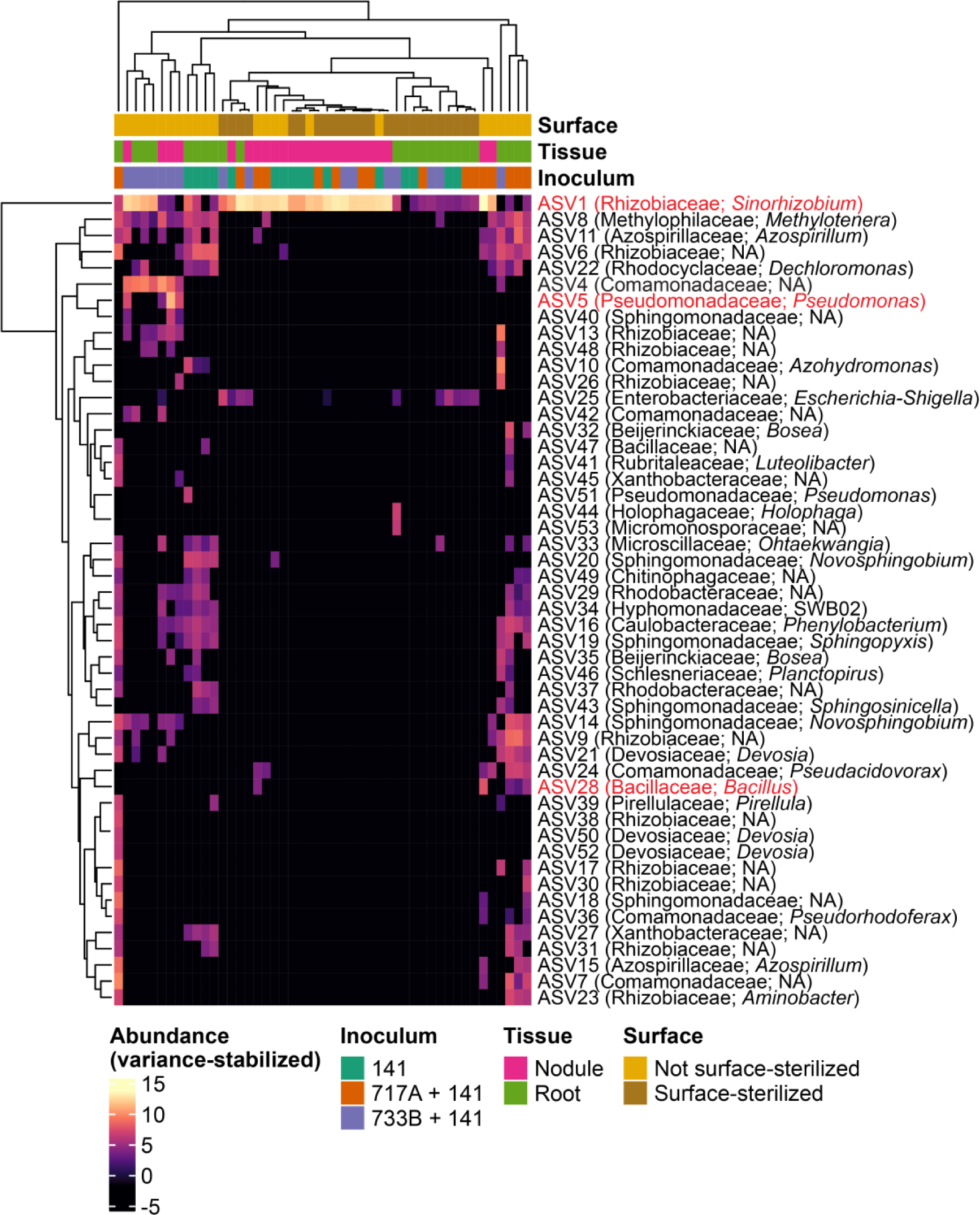
Amplicon sequence variants inferred from nodule and root samples of *Medicago truncatula*. Amplicon sequence variant (ASV) counts transformed by variance-stabilization are given in the heatmap matrix. Columns correspond to individual DNA libraries sequenced and extraced from *M. truncatula* samples across a combination of inoculation treatments, tissue sections, and surface-sterilization treatments. Rows represent the abundance of individual ASVs in a sample. For simplicity, only the 50 most abundant ASVs are shown. Family and genus taxonomy labels are as obtained by DECIPHER (Wright 2016) using the SILVA SSU r138 (Quast et al. 2012) training set, NA: unassigned. AVS highlighted in red are ASVs of interest suspected to represent our inoculum strains.

Our goal was to determine whether we could recover our inoculum strains from nodules in the expected treatments, which would delineate these strains as either nodule endophytes or tissue surface epiphytes. After aligning our inferred ASVs against the 16S rRNA sequences of our isolate genomes, we found that ASV1 and ASV5 represent our *Sinorhizobium* and *Pseudomonas* sp. 733B inocula, respectively (Supplemental Results, Table S9). ASV1 (*Sinorhizobium* based on SILVA references) was abundant in every inoculum treatment, tissue section, and surface-sterilization treatment. The abundance of ASV1 was not different between samples inoculated with *Sinorhizobium* alone and either 717A + 141 or 733B + 141 samples (*P* > 0.05, Wald test, FDR adjusted) (Figure 4, Figure S1A-B, Table S12). As expected, ASV1 was significantly more abundant among nodule samples compared to root samples (*P* < 0.01, Wald test, FDR adjusted) (Figure S1C). Thus we find no evidence that *Bacillus* sp. 717A or *Pseudomonas* sp. 733B influence the abundance of *Sinorhizobium* once they are within the nodules. ASV5 was assigned to *Pseudomonas* SILVA reference sequences and was only detected in samples from *Pseudomonas* sp. 733B co-inoculated plants. This ASV was present in non-surface-sterilized nodule samples plus one additional surface-sterilized root sample at trace levels, but was significantly more abundant in nodule samples compared to root samples (*P* < 0.01, Wald test, FDR adjusted) (Figure 4, Figure S1C, Table S12). For *Bacillus* sp. 717A, we determined that ASV28 is the most likely representative of out of all other ASVs inferred from the amplicon data. This ASV was found only in non-surface-sterilized nodule and non-surface-sterilized root samples from *Bacillus* sp. 717A co-inoculated plants, with no difference in abundance between the two tissue sections (*P* > 0.05, Wald test, FDR adjusted) (Figure 4, Figure S1A, Table S12).

To determine whether inoculum strains were differentially abundant across the *M. truncatula* rhizosphere versus the endosphere, we compared surface-sterilized to non-surface-sterilized samples (Figure S1D-E). Strains that are endosphere-associated should either be enriched in the surface-sterilized groups (indicating preference for the endosphere over the rhizosphere) or not differentially abundant between the two groups (indicating similar levels across the endosphere and rhizosphere). However, strains that are rhizosphere-associated should be enriched in non-surface-sterilized groups. In the nodule samples, ASV5 [*Pseudomonas* sp. 733B] and ASV28 [*Bacillus* sp. 717A] were enriched in non-surface-sterilized samples (*P* < 0.05, *P* < 0.01, Wald test, FDR adjusted), while ASV1 [*Sinorhizobium*] was not differentially abundant (*P* > 0.05, Wald test, FDR adjusted) (Figure S1E, Table S12). In the root samples, ASV28 was enriched in non-surface-sterilized samples (*P* < 0.01, Wald test, FDR adjusted). Although ASV1 was more abundant in surface-sterilized root samples compared to non-surface-sterilized root samples and ASV5 was less abundant (*P* < 0.05, Wald test, FDR adjusted), neither ASV met our minimum fold-change criterion of 1.5 to be considered enriched in either surface-sterilized or non-surface-sterilized root samples (Figure S1F, Table S12). Collectively, these comparisons show that *Sinorhizobium* was indeed endosphere-associated in both root and nodule sections. *Bacillus* sp. 717A, on the other hand, was rhizosphere associated in nodules and root sections, while *Pseudomonas* sp. 733B was rhizosphere associated in nodules and endosphere-associated in root sections.

### Sinorhizobium, Bacillus, and Pseudomonas associations beyond this study

To further investigate the ecological relevance of our NREs, we asked whether our three ASVs of interest, representing the three inoculum strains, could also be detected among amplicon data from a separate study of the leaf, root, and nodule endophytes of *M. truncatula* endophytes grown in native field soil (Brown et al. 2020). Aligning our ASVs of interest against the 50 most abundant sequences generated by Brown et al. (2020) revealed imperfect but numerous alignments to taxa in their study (Table S10). The best alignment of ASV1 was OTU00001, an “*Ensifer*” bacterium (now reclassified as *Sinorhizobium*) that was found to be a significant nodule endophyte in the Brown et al. (2020) study, at 99% identity with 92% coverage of the OTU (Table S10). Our ASV28 [*Bacillus* sp. 717A] aligned to OTU00009 at 100% identity and 92% coverage (Table S10). This OTU was a significant rhizosphere member in Brown et al. (2020) and was labelled as a “*Paenisporosarcina*” in the order Bacillales, inferred by aligning OTUs against a SILVA 16S reference alignment (v123). Lastly, our ASV5 [*Pseudomonas* sp. 733B] aligned equally well to OTU00013 and OTU00003, which were labeled by Brown et al. (2020) as “*Streptomyces*” and “*Pseudomonas*”, respectively, at 99% identity and 92% coverage (Table S10). Because the OTU sequence lengths in Brown et al. (2020) were over 100 bases shorter than the ASVs generated here, the alignments of ASV5 to both *Streptomyces* and *Pseudomonas* OTUs are likely due to the lower sequence resolution in their study. Given that ASV5 [*Pseudomonas* sp. 733B] aligned perfectly to a region of the 16S rRNA gene in our *Pseudomonas* sp. 733B genome assembly, we believe that our ASV5 [*Pseudomonas* sp. 733B] is likely closely related to *Pseudomonas* OTU00003, a significant member of the leaf phyllosphere in Brown et al. (2020).

## DISCUSSION

Given the ecological and economic importance of legume productivity in diverse biotic contexts (Sprent 1987; Herridge et al. 2008a), here we asked how the benefits of legume-rhizobium symbiosis (shoot mass for the host, number of nodules for the rhizobium) changed for both partners due to the presence of non-rhizobial endophytes (NREs) that share the nodule alongside rhizobia. Using four NRE strains, we inferred their taxonomy, re-identified the NREs from root and/or nodule tissues, and showed that two of the four NREs we examined reduced the benefits both legumes and rhizobia receive from the interaction. We discuss the potential mechanisms for, and implications of, these main results below.

While it is infeasible to capture the effects of all surrounding taxa on legume-rhizobium symbiosis, investigating interactions with microbes present inside root nodules is an important step towards understanding legume-rhizobium symbiosis in its full context. Several studies of legume-rhizobium symbiosis have demonstrated how genetic variation in either of these partners can influence fitness outcomes, in addition to interactions with other plants and microbes (Heath and Tiffin 2009; Brown et al. 2020; Batstone et al. 2022a). For example, one study found that variation in host control among six lines of *Acmispon strigosus* influenced the variation in symbiont effectiveness among *Bradyrhizobium* populations (Wendlandt et al. 2019). Far fewer studies have explicitly considered genetic variation in other microbial constituents present in the soil community (Tsiknia et al. 2020). We show that genetic variation within NREs (*Pseudomonas* spp. 702A vs. 733B and *Paenibacillus* sp. 522 vs. *Bacillus* sp. 717A) leads to variation in fitness-related traits of both the *M. truncatula* host and the *S. meliloti* symbiont. Our results suggest that NREs present in nodules alongside rhizobia can have indirect effects that must be considered if the outcomes of the legume-rhizobium symbiosis in real soil communities are to be predicted more accurately.

### Effects on plant fitness

Symbionts in a microbiome do not always evolve to benefit their plant host, as microbes have their own fitness interest that do not necessarily align with host fitness (Klein et al. 2021). In contrast to previous studies, the NREs included here were isolated from the same region as our focal *Sinorhizobium* strain and negatively impacted the legume-rhizobium symbiosis by decreasing the number of nodules formed and shoot mass. The vast majority of NREs described to date have demonstrated positive or neutral effects on plant or rhizobium performance by increasing host shoot mass or the number of nodules formed (Khan, 2019; Martínez-Hidalgo & Hirsch, 2017). Since two NREs in this study (*Bacillus* sp. 717A and *Pseudomonas* sp. 733B) negatively impacted the legume-rhizobium mutualism while the other two (*Paenibacillus* sp. 522 and *Pseudomonas* sp. 702A) showed neutral effects, NREs should not generally be considered beneficial. Our experimental approach allowed us to examine the impacts co-inoculation with NREs have on plant growth. But we recognize that NREs might possess other traits from which plants could benefit, including drought resistance, nutrient acquisition, and herbivory resistance. Further studies that examine rhizobia-NRE interactions in the context of a complex soil microbiome under various environmental conditions would be fruitful for understanding the full range of potential effects of NREs on host fitness.

The negative effects of *Bacillus* sp. 717A and *Pseudomonas* sp. 733B on plant growth could have been due to several non-mutually exclusive mechanisms, including that these strains: i) represent plant pathogens, whereby plants perform worse in the presence of these NREs compared to when they are not associated with them regardless of the presence of rhizobia, ii) reduce *Sinorhizobium*’s ability to infect and form nodules on plants; and/or iii) reduce *Sinorhizobium*’s ability to proliferate and/or fix N within nodules. The results from our nitrogen addition experiment, whereby plants were provided with sufficient levels of N critical for minimal growth (Küster 2013), provided little evidence for i), that NREs are plant pathogens; plants performed similarly when inoculated by NREs compared to sterile media. Our results also do not support iii), that NREs impede *Sinorhizobium*’s ability to inhabit and proliferate within nodules because we found little evidence for changes in *Sinorhizobium* ASV abundance within nodules due to NRE co-inoculation. Instead, our results are most consistent with ii), that NREs interfere with *Sinorhizobium*’s ability to form nodules on host plants, given the significant reduction in nodules formed by plants when co-inoculated with NREs compared to *Sinorhizobium* alone.

Finding a reduction in nodulation despite plants being N starved in our greenhouse experiment suggests that these NREs inhibited nodulation, negatively impacting N acquisition by the host from rhizobial partners. When the only source of N is that fixed by rhizobia, legumes will continue to form nodules until they acquire sufficient levels of N via a tightly regulated process (Bauer 1981; Caetano-Anollés and Gresshoff 1991). A recent study in which legumes were either co-inoculated with two strains of rhizobia or singly inoculated with each strain individually found a similar reduction in nodulation due to interference competition between strains, whereby the growth of each strain was inhibited by the presence of the other (Rahman et al. 2023). Given their ability to co-inhabit root and nodule tissues, direct interactions between rhizobia and NREs are likely. In other studies, rhizobia-NRE interactions were found to impact nodulation in a variety of ways. For example, *S. meliloti* was found to cross-utilize siderophores produced by *Exiguobacterium* NREs in co-inoculations, which stimulated nodule formation and increased the mass of their *Trigonella foenum-graecum* host (Rajendran et al. 2012). Additionally, production of indole-3-acetic acid by *Pseudomonas trivialis* 3Re27 and *Pseudomonas extremorientalis* TSAU20 stimulated nodule formation by *Rhizobium galegae* on roots of *Galega orientalis* and increased *G. orientalis* growth (Egamberdieva et al. 2010). Furthermore, *Bacillus subtilis* UD1022 was found to antagonistically downregulate quorum sensing and biofilm formation by *Sinorhizobium meliloti* Rm8530, which is critical to initiate root nodule formation (Rosier et al. 2021). Future studies could determine whether the NREs *Pseudomonas sp.* 733B and *Bacillus sp.* 717A act antagonistically against *Sinorhizobium* via extracellular secretions that inhibit nodule formation, or whether they compete with *Sinorhizobium* in root tissues. Either or both of these mechanisms could explain the indirect impacts of these NREs on plant growth. Overall, our results highlight that NREs have the potential to negatively impact plants *indirectly* through their interactions with rhizobia, which in turn, impacts the fitness benefits received by both legumes and rhizobia. Improving symbiotic outcomes in natural or managed fields will therefore require examining the community context in which the symbiosis occurs rather than the focal partners alone.

### Cohabitation of Sinorhizobium and NREs

Because they shift the costs and benefit exchange, biotic players such as NREs may have important roles in shaping mutualism evolution (Strauss 1991; Strauss and Irwin 2004; Keller et al. 2018). Evolutionary theory predicts that mutualisms are susceptible to “cheaters” that reap rewards from their symbionts without paying any costs (Jones et al. 2015). One current objective of coevolutionary research is to understand how genetic variation and abiotic factors stabilize or destabilize mutualisms (Sachs et al. 2004; Heath and Tiffin 2009; Jones et al. 2015; Batstone et al. 2018, 2022b). Some perspectives suggest that maintaining variation in partner quality, even continuing to associate with potential cheaters, offers a selective advantage to the host if the benefits obtained depend on environmental conditions (Batstone et al. 2018). By extension, the benefits exchanged in symbiosis are likely to depend on the biotic context, and more specifically, the microbiome in which the focal symbiosis unfolds. The natural co-habitation of rhizobia and NREs that impact legume-rhizobia symbiosis here could provide yet another mechanism for maintaining variation in partner quality, if quality depends on the identities of the NREs present within a microbiome (Batstone et al. 2018).

Soil bacteria interact with each other and influence the fitness of their host plant with and without infection (de la Fuente Cantó et al. 2020). Whether the microbes behind these interactions inhabit the nodule/root endosphere or rhizosphere has different implications for community composition and the phenotypic outcomes of symbioses (Brown et al. 2020). While the NRE strains used here were originally isolated from cultures of the *M. truncatula* nodule endosphere, we were not able to confirm reinfection of nodule endosphere by either NRE strain, although we did detect both on surface nodule and root tissues and *Pseudomonas* sp. 733B in the root endosphere at relatively low levels. Our findings do not rule-out that these NREs occupied nodules alongside *Sinorhizobium*. Low-abundance bacteria can have significant impacts on the surrounding community (Lynch and Neufeld 2015; Jousset et al. 2017). These community members may be difficult to detect in amplicon sequencing data if their sequence counts in a sample are too low compared to other community members (Huse et al. 2010; Paulson et al. 2013; Escudié et al. 2018). Given *Sinorhizobium*’s extensive population growth and genome duplication within nodules (Mergaert et al. 2006), sequencing technologies that enrich for particular targets, such as capture sequencing (Hayden et al. 2022), may be required to detect low frequency non-rhizobial nodule occupants.

### Identifying potential Pseudomonas and Bacillus NREs elsewhere

The *Pseudomonas* and *Bacillus* NRE strains studied here were originally isolated from nodules of *M. truncatula* grown in soils collected from its natural range. By comparing our sequencing results to those of another study that grew *M. truncatula* in such soils (Brown et al. 2020), we found OTUs representing taxa closely related to *Bacillus* sp. 717A and *Pseudomonas* sp. 733B to be among the most abundant community members. This relationship suggests that NREs in this study are likely reflective of NREs in the natural microbiome of *M. truncatula*. This presents opportunities to understand the mechanisms by which complex root-associated communities establish and influence each other using this as a model multi-player symbiosis.

### Concluding remarks

Here we show that some NREs can reduce the benefits of legume-rhizobium symbiosis. In our study, *Paenibacillus* sp. 522, *Bacillus* sp. 717A, as well as *Pseudomonas* spp. 702A and 733B are each phylogenetically distinct strains representing four novel species without high nucleotide similarity to existing RefSeq representatives. When co-inoculated alongside *S. meliloti*, two of these strains (717A and 733B) were re-identified on root and nodule tissues of *M. truncatula* and were found to inhibit nodulation and plant growth. We have little evidence that NREs impacted *Sinorhizobium*’s ability to proliferate within nodules, whereas we show stronger support that NREs negatively impacted *Sinorhizobium*’s ability to form nodules. Thus, it is likely that the NREs and *Sinorhizobium* exhibited competitive interactions with rhizobia outside the roots or in the soil. Our results highlight that intraspecific variation within NREs can generate variable fitness outcomes for both partners in the legume-rhizobium symbiosis, meaning that predicting the impact of these “off-target” strains have on legume-rhizobium symbiosis will require more than just testing a single representative strain across different species. Thankfully, with the rise of more affordable sequencing technologies, the variation in third-party species such as the NREs identified here can be uncovered more feasibly. Coupling sequencing with experiments to test the potential indirect effects of NREs present in natural or managed soil microbiomes will facilitate a better understanding of the coevolutionary dynamics in complex microbiome communities.

## ACKNOWLEDGMENTS

The authors thank all current and past members of the Heath Lab at the University of Illinois at Urbana-Champaign who have assisted with and provided intellectual support for the work presented here. In particular, the authors thank Michael Grillo and Alex Riley for their initial efforts in isolating and identifying the bacterial strains used in this study. The authors also thank Karla Griesbaum, Nevers Mushimata, and Riley Popp for assisting with greenhouse preparations, plant harvests, and nodule dissections. The authors additionally thank Mario Cerón Romero for advice on bioinformatics analyses as well as Hunter Cobbley, David Vereau Gorbitz, and Ivan Sosa Marquez for assistance with DNA extractions and genome sequencing. We would also like to thank members of McMaster Biology’s Data Lunch run by Dr. Ben Bolker for their invaluable advice on the statistical approach we implemented.

## AUTHOR CONTRIBUTION STATEMENT

Conceptualization, J.C.K., R.B.D., and K.D.H.; Methodology, J.C.K., R.B.D., and K.D.H.; Software, J.C.K. and R.B.D.; Validation, J.C.K., R.B.D., and K.D.H.; Formal Analysis, J.C.K. and R.B.D.; Investigation, J.C.K., R.B.D., and K.D.H.; Resources, K.D.H.; Data curation, J.C.K., R.B.D., and K.D.H.; Writing — Original Draft, J.C.K.; Writing — Review & Editing, J.C.K., R.B.D., and K.D.H.; Visualization, J.C.K. and R.B.D.; Supervision, R.B.D. and K.D.H.; Project Administration, K.D.H. and R.B.D.; Funding Acquisition, K.D.H.

## DATA AVAILABILITY STATEMENT

All scripts and intermediate files are available at github.com/jamesck2/coinoculation-endophytes. The raw 16S V3-V4 amplicon reads analyzed here are available at NCBI BioProject PRJNA1077341 and the whole-genome sequences of *Sinorhizobium meliloti* 141, *Bacillus* sp. 717A, and *Pseudomonas* sp. 733B are available at NCBI BioProject PRJNA1077595.

## SUPPLEMENTARY FIGURES

**Figure S1.**
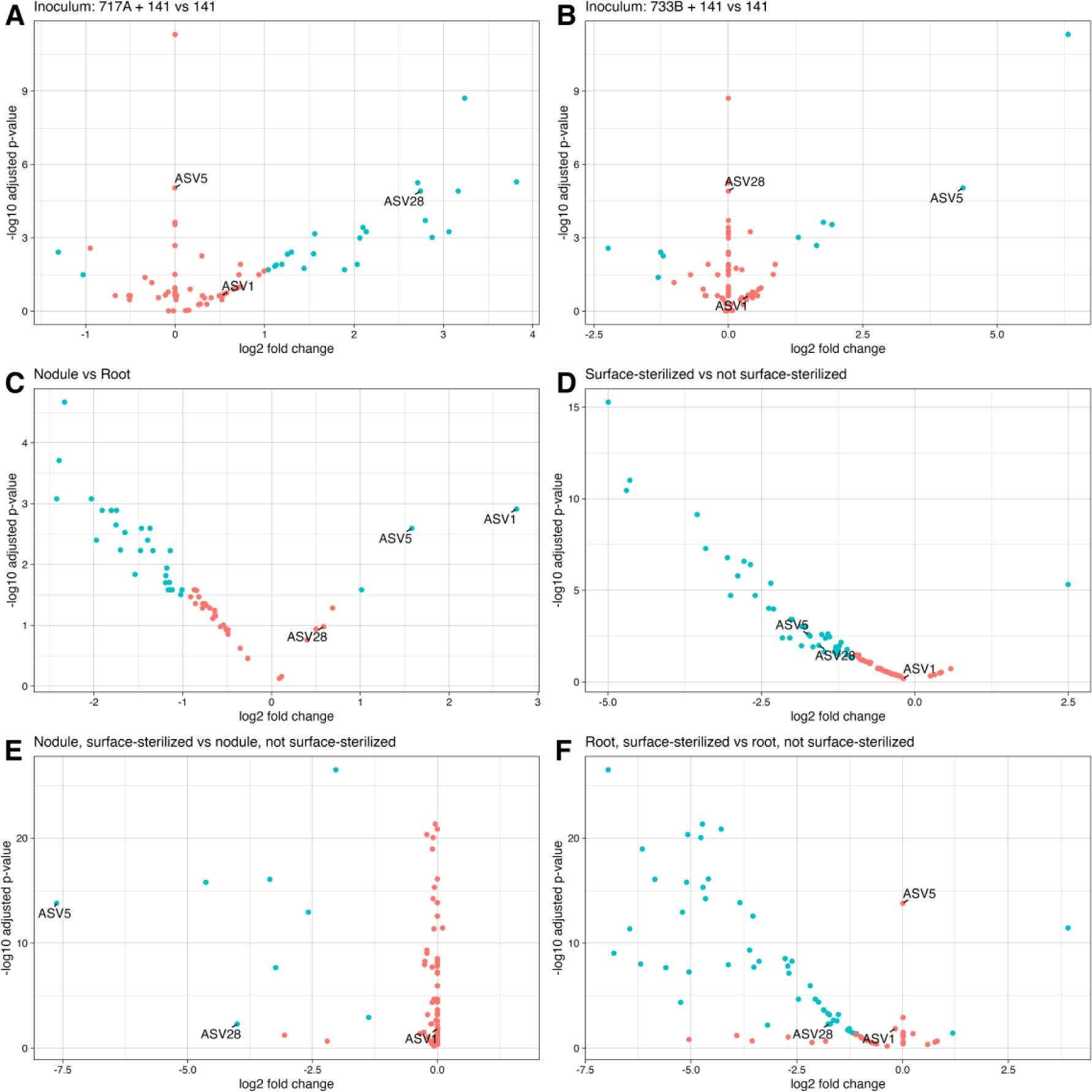
Differential abundance of ASVs across inocula, tissue sections, and surface-sterilization treatments. Using a minimum fold change 1.5 and *P* (false-discovery rate adjusted) < 0.05, amplicon sequence variants (ASVs) diferentially abundant across contrasts of inocula, tissues, and surface-sterilization treatments were inferred. Points refer to individual ASVs, blue points correspond to signifiicant ASVs while red points are insignificant. The second term in plot titles indicate baselines in comparisons. ASVs representing inocula strains are labeled, ASV1 = *S. meliloti* 141; ASV28 = *Bacillus* sp. 717A; ASV5 = *Pseudomonas* sp. 733B. (A) ASV28 was enriched in *Bacillus* sp. 717A + *S. meliloti* 141 co-inoculated samples compared to samples inoculated with *S. meliloti* 141 only. Similarly, (B) ASV5 was enriched in *Pseudomonas* sp. 733B + *S. meliloti* 141 co-inoculated samples compared to samples inoculated with *S. meliloti* 141 only. (C) Both ASV5 and ASV1 were enriched in nodule samples compared to root samples. (D) ASV5 and ASV28 were enriched in non-surface-sterilized samples compared to surface-sterilized samples, while ASV1 was not differentially abundant across either group. (E) Among just nodule samples, ASV5 and ASV28 were enriched in non-surface-sterilized samples while ASV1 was not differentially abundant. (F) Among just root samples, only ASV28 was enriched in non-surface-sterilized root samples while ASV1 and ASV5 were not different across either group.

## SUPPLEMENTARY TABLE LEGENDS

**Table S1.** Type strain genome information for the TYGS (Meier-Kolthoff and Göker 2019) phylogenomic analyses.

**Table S2.** Pairwise intergenomic distance calculations between all user and type strains performed by TYGS (Meier-Kolthoff and Göker 2019), as used in phylogenomic tree construction.

**Table S3.** Raw phenotypic data collected from the indirect effects greenhouse experiment.

**Table S4.** Pairwise contrasts of estimated marginal means (Lenth, 2022) with Dunnet-adjusted *P*-valuesClick or tap here to enter text. for shoot mass and nodule number for between inoculation treatments in the indirect effects greenhouse experiment.

**Table S5.** Estimated marginal means (Lenth, 2022) for shoot mass and nodule number for each inoculation treatment in the indirect effects greenhouse experiment.

**Table S6.** Raw phenotypic data collected from the direct effects nitrogen-addition experiment.

**Table S7.** Amplicon sequence variant (ASV) table generated by DADA2 (Callahan et al. 2016) from amplified 16S V3-V4 rRNA regions of DNA extracted from nodule and root tissues of co-inoculated and single-inoculated *M. truncatula* plants. Column names refer to ASV names, row names are individual DNA samples. For sample metadata, see Table S11.

**Table S8.** Nucleotide sequences of ASVs present in Table S7.

**Table S9.** Nucleotide BLAST (Camacho et al. 2009) of ASV sequences in Table S7 against 16S rRNA gene annotations from *Sinorhizobium meliloti* 141, *Bacillus* sp. 717A, and *Pseudomonas* sp. 733B. Minimum percent identity of 80% and maximum e-value of 0.01.

**Table S10.** Nucleotide BLAST (Camacho et al. 2009) of ASV sequences in Table S7 against the top 50 OTU sequences from Brown et al. (Brown et al. 2020). Minimum percent identity of 80% and maximum e-value of 0.01.

**Table S11.** Metadata for tissue occupancy experiment DNA samples as reported in Table S7.

**Table S12.** Wald test results from DESeq2 (Love et al. 2014) comparing the difference in abundance for ASVs present in Table S7 across inoculum treatments, tissue sections, and surface-sterilization treatments.

## REFERENCES

Afkhami, M.E., Almeida, B.K., Hernandez, D.J., Kiesewetter, K.N., and Revillini, D.P. 2020. Tripartite mutualisms as models for understanding plant–microbial interactions. Curr Opin Plant Biol 56: 28–36. doi:10.1016/j.pbi.2020.02.003.

Bakker, M.G., Schlatter, D.C., Otto-Hanson, L., and Kinkel, L.L. 2014. Diffuse symbioses: roles of plant–plant, plant–microbe and microbe–microbe interactions in structuring the soil microbiome. Mol Ecol 23(6): 1571–1583. doi:10.1111/mec.12571.

Batstone, R.T., Burghardt, L.T., and Heath, K.D. 2022a. Phenotypic and genomic signatures of interspecies cooperation and conflict in naturally occurring isolates of a model plant symbiont. Proceedings of the Royal Society B: Biological Sciences 289(1978). Royal Society Publishing. doi:10.1098/rspb.2022.0477.

Batstone, R.T., Carscadden, K.A., Afkhami, M.E., and Frederickson, M.E. 2018. Using niche breadth theory to explain generalization in mutualisms. Ecology 99(5): 1039– 1050.

Batstone, R.T., Lindgren, H., Allsup, C.M., Goralka, L.A., Riley, A.B., Grillo, M.A., Marshall-Colon, A., and Heath, K.D. 2022b. Genome-Wide Association Studies across Environmental and Genetic Contexts Reveal Complex Genetic Architecture of Symbiotic Extended Phenotypes. mBio 13(6). American Society for Microbiology. doi:10.1128/mbio.01823-22.

Bauer, W.D. 1981. Infection of Legumes by Rhizobia. Annu Rev Plant Physiol 32(1): 407–449. doi:10.1146/annurev.pp.32.060181.002203.

Benjamini, Y., and Hochberg, Y. 1995. Controlling the False Discovery Rate: A Practical and Powerful Approach to Multiple Testing. Journal of the Royal Statistical Society: Series B (Methodological) 57(1): 289–300. doi:10.1111/j.2517-6161.1995.tb02031.x.

Bolin, L.G., Lennon, J.T., and Lau, J.A. 2023. Traits of soil bacteria predict plant responses to soil moisture. Ecology 104(2). doi:10.1002/ecy.3893.

Brown, S., Grillo, M., Podowski, J., and Heath, K.D. 2020. Soil origin and plant genotype structure distinct microbiome compartments in the model legume Medicago truncatula. Microbiome 8(139). doi:10.21203/rs.3.rs-21175/v1.

Caetano-Anollés, G., and Gresshoff, P.M. 1991. Plant genetic control of nodulation. Annu Rev Microbiol 45(1): 345–382. doi:10.1146/annurev.mi.45.100191.002021.

Callahan, B.J., McMurdie, P.J., Rosen, M.J., Han, A.W., Johnson, A.J.A., and Holmes, S.P. 2016. DADA2: High-resolution sample inference from Illumina amplicon data. Nat Methods 13(7): 581–583. doi:10.1038/nmeth.3869.

Camacho, C., Coulouris, G., Avagyan, V., Ma, N., Papadopoulos, J., Bealer, K., and Madden, T.L. 2009. BLAST+: architecture and applications. BMC Bioinformatics 10(1): 421. doi:10.1186/1471-2105-10-421.

Egamberdieva, D., Berg, G., Lindström, K., and Räsänen, L.A. 2010. Co-inoculation of Pseudomonas spp. with Rhizobium improves growth and symbiotic performance of fodder galega (Galega orientalis Lam.). Eur J Soil Biol 46(3–4): 269–272. doi:10.1016/j.ejsobi.2010.01.005.

Escudié, F., Auer, L., Bernard, M., Mariadassou, M., Cauquil, L., Vidal, K., Maman, S., Hernandez-Raquet, G., Combes, S., and Pascal, G. 2018. FROGS: Find, Rapidly, OTUs with Galaxy Solution. Bioinformatics 34(8): 1287–1294. doi:10.1093/bioinformatics/btx791.

Fox, S.L., O’Hara, G.W., and Bräu, L. 2011. Enhanced nodulation and symbiotic effectiveness of Medicago truncatula when co-inoculated with Pseudomonas fluorescens WSM3457 and Ensifer (Sinorhizobium) medicae WSM419. Plant Soil 348(1–2): 245–254. doi:10.1007/s11104-011-0959-8.

Garnier, S., Ross, N., Rudis, R., Camargo, A.P., Sciaini, M., and Scherer, C. 2023. viridis(Lite) - Colorblind-Friendly Color Maps for R.

Haney, C.H., Samuel, B.S., Bush, J., and Ausubel, F.M. 2015. Associations with rhizosphere bacteria can confer an adaptive advantage to plants. Nat Plants 1(6): 15051. doi:10.1038/nplants.2015.51.

Hayden, H.S., Joshi, S., Radey, M.C., Vo, A.T., Forsberg, C., Morgan, S.J., Waalkes, A., Holmes, E.A., Klee, S.M., Emond, M.J., Singh, P.K., and Salipante, S.J. 2022. Genome Capture Sequencing Selectively Enriches Bacterial DNA and Enables Genome-Wide Measurement of Intrastrain Genetic Diversity in Human Infections. mBio 13(5). doi:10.1128/mbio.01424-22.

Heath, K.D., and Tiffin, P. 2007. Context dependence in the coevolution of plant and rhizobial mutualists. Proceedings of the Royal Society B: Biological Sciences 274(1620): 1905–1912. Royal Society. doi:10.1098/rspb.2007.0495.

Heath, K.D., and Tiffin, P. 2009. Stabilizing mechanisms in a legume-rhizobium mutualism. Evolution (N Y) 63(3): 652–662. doi:10.1111/j.1558-5646.2008.00582.x.

Herridge, D.F., Peoples, M.B., and Boddey, R.M. 2008a. Global inputs of biological nitrogen fixation in agricultural systems. Plant Soil 311(1–2): 1–18. doi:10.1007/s11104-008-9668-3.

Herridge, D.F., Peoples, M.B., and Boddey, R.M. 2008b. Global inputs of biological nitrogen fixation in agricultural systems. Plant Soil 311(1–2): 1–18. doi:10.1007/s11104-008-9668-3.

Huse, S.M., Welch, D.M., Morrison, H.G., and Sogin, M.L. 2010. Ironing out the wrinkles in the rare biosphere through improved OTU clustering. Environ Microbiol 12(7): 1889–1898. doi:10.1111/j.1462-2920.2010.02193.x.

Johnson, N.C., Graham, J., and Smith, F.A. 1997. Functioning of mycorrhizal associations along the mutualism–parasitism continuum*. New Phytologist 135(4): 575–585. doi:10.1046/j.1469-8137.1997.00729.x.

Jones, E.I., Afkhami, M.E., Akçay, E., Bronstein, J.L., Bshary, R., Frederickson, M.E., Heath, K.D., Hoeksema, J.D., Ness, J.H., Pankey, M.S., Porter, S.S., Sachs, J.L., Scharnagl, K., and Friesen, M.L. 2015, November 1. Cheaters must prosper: Reconciling theoretical and empirical perspectives on cheating in mutualism. Blackwell Publishing Ltd. doi:10.1111/ele.12507.

Keller, K.R., Carabajal, S., Navarro, F., and Lau, J.A. 2018. Effects of multiple mutualists on plants and their associated arthropod communities. Oecologia 186(1): 185–194. doi:10.1007/s00442-017-3984-3.

Kent, A.D., and Triplett, E.W. 2002. Microbial Communities and Their Interactions in Soil and Rhizosphere Ecosystems. Annu Rev Microbiol 56(1): 211–236. doi:10.1146/annurev.micro.56.012302.161120.

Klein, M., Stewart, J.D., Porter, S.S., Weedon, J.T., and Kiers, E.T. 2021. Evolution of manipulative microbial behaviors in the rhizosphere. John Wiley and Sons Inc. doi:10.1111/eva.13333.

Klein, M., Stewart, J.D., Porter, S.S., Weedon, J.T., and Kiers, E.T. 2022. Evolution of manipulative microbial behaviors in the rhizosphere. Evol Appl 15(10): 1521–1536. doi:10.1111/eva.13333.

Kolde, R. 2019. pheatmap: Pretty Heatmaps.

Küster, H. 2013. Medicago truncatula. In Brenner’s Encyclopedia of Genetics: Second Edition. Elsevier Inc. pp. 335–337. doi:10.1016/B978-0-12-374984-0.00915-3.

Kuzmanović, N., Fagorzi, C., Mengoni, A., Lassalle, F., and diCenzo, G.C. 2022. Taxonomy of Rhizobiaceae revisited: proposal of a new framework for genus delimitation. Int J Syst Evol Microbiol 72(3). doi:10.1099/ijsem.0.005243.

Kuznetsova, A., Brockhoff, P.B., and Christensen, R.H.B. 2017. lmerTest Package: Tests in Linear Mixed Effects Models. J Stat Softw 82(13). doi:10.18637/jss.v082.i13.

de la Fuente Cantó, C., Simonin, M., King, E., Moulin, L., Bennett, M.J., Castrillo, G., and Laplaze, L. 2020. An extended root phenotype: the rhizosphere, its formation and impacts on plant fitness. The Plant Journal 103(3): 951–964. doi:10.1111/tpj.14781.

Laczny, C.C., Sternal, T., Plugaru, V., Gawron, P., Atashpendar, A., Margossian, H.H., Coronado, S., der Maaten, L. van, Vlassis, N., and Wilmes, P. 2015. VizBin - an application for reference-independent visualization and human-augmented binning of metagenomic data. Microbiome 3(1): 1. doi:10.1186/s40168-014-0066-1.

Lau, J.A., and Lennon, J.T. 2012. Rapid responses of soil microorganisms improve plant fitness in novel environments. Proceedings of the National Academy of Sciences 109(35): 14058–14062. doi:10.1073/pnas.1202319109.

Lefort, V., Desper, R., and Gascuel, O. 2015. FastME 2.0: A Comprehensive, Accurate, and Fast Distance-Based Phylogeny Inference Program: Table 1. Mol Biol Evol 32(10): 2798–2800. doi:10.1093/molbev/msv150.

Lenth, R. v. 2022. emmeans: Estimated Marginal Means, aka Least-Squares Means. Available from https://CRAN.R-project.org/package=emmeans.

Love, M.I., Huber, W., and Anders, S. 2014. Moderated estimation of fold change and dispersion for RNA-seq data with DESeq2. Genome Biol 15(12): 550. doi:10.1186/s13059-014-0550-8.

Martínez-Hidalgo, P., Galindo-Villardón, P., Igual, J.M., and Martínez-Molina, E. 2014. Micromonospora from nitrogen fixing nodules of alfalfa (Medicago sativa L.). A new promising Plant Probiotic Bacteria. Sci Rep 4. Nature Publishing Group. doi:10.1038/srep06389.

Martínez-Hidalgo, P., and Hirsch, A.M. 2017. The nodule microbiome: N2fixing rhizobia do not live alone. American Phytopathological Society. doi:10.1094/PBIOMES-12-16-0019-RVW.

McMurdie, P.J., and Holmes, S. 2013. phyloseq: An R Package for Reproducible Interactive Analysis and Graphics of Microbiome Census Data. PLoS One 8(4): e61217. doi:10.1371/journal.pone.0061217.

Meier-Kolthoff, J.P., Auch, A.F., Klenk, H.-P., and Göker, M. 2013. Genome sequence-based species delimitation with confidence intervals and improved distance functions. BMC Bioinformatics 14(1): 60. doi:10.1186/1471-2105-14-60.

Meier-Kolthoff, J.P., Carbasse, J.S., Peinado-Olarte, R.L., and Göker, M. 2022. TYGS and LPSN: a database tandem for fast and reliable genome-based classification and nomenclature of prokaryotes. Nucleic Acids Res 50(D1): D801–D807. doi:10.1093/nar/gkab902.

Meier-Kolthoff, J.P., and Göker, M. 2019. TYGS is an automated high-throughput platform for state-of-the-art genome-based taxonomy. Nat Commun 10(1): 2182. doi:10.1038/s41467-019-10210-3.

Meier-Kolthoff, J.P., Hahnke, R.L., Petersen, J., Scheuner, C., Michael, V., Fiebig, A., Rohde, C., Rohde, M., Fartmann, B., Goodwin, L.A., Chertkov, O., Reddy, T., Pati, A., Ivanova, N.N., Markowitz, V., Kyrpides, N.C., Woyke, T., Göker, M., and Klenk, H.-P. 2014. Complete genome sequence of DSM 30083T, the type strain (U5/41T) of Escherichia coli, and a proposal for delineating subspecies in microbial taxonomy. Stand Genomic Sci 9(1): 2. doi:10.1186/1944-3277-9-2.

Mergaert, P., Uchiumi, T., Alunni, B., Evanno, G., Cheron, A., Catrice, O., Mausset, A.-E., Barloy-Hubler, F., Galibert, F., Kondorosi, A., and Kondorosi, E. 2006. Eukaryotic control on bacterial cell cycle and differentiation in the *Rhizobium* – legume symbiosis. Proceedings of the National Academy of Sciences 103(13): 5230–5235. doi:10.1073/pnas.0600912103.

Neuwirth, E. 2022. RColorBrewer: ColorBrewer Palettes.

Nikodemova, M., Holzhausen, E.A., Deblois, C.L., Barnet, J.H., Peppard, P.E., Suen, G., and Malecki, K.M. 2023. The effect of low-abundance OTU filtering methods on the reliability and variability of microbial composition assessed by 16S rRNA amplicon sequencing. Front Cell Infect Microbiol 13. doi:10.3389/fcimb.2023.1165295.

O’Brien, A.M., Jack, C.N., Friesen, M.L., and Frederickson, M.E. 2021. Whose trait is it anyways? Coevolution of joint phenotypes and genetic architecture in mutualisms. Proceedings of the Royal Society B: Biological Sciences 288(1942): 20202483. doi:10.1098/rspb.2020.2483.

Paulson, J.N., Stine, O.C., Bravo, H.C., and Pop, M. 2013. Differential abundance analysis for microbial marker-gene surveys. Nat Methods 10(12): 1200–1202. doi:10.1038/nmeth.2658.

Porter, S.S., Chang, P.L., Conow, C.A., Dunham, J.P., and Friesen, M.L. 2017. Association mapping reveals novel serpentine adaptation gene clusters in a population of symbiotic Mesorhizobium. ISME J 11(1): 248–262. doi:10.1038/ismej.2016.88.

Quast, C., Pruesse, E., Yilmaz, P., Gerken, J., Schweer, T., Yarza, P., Peplies, J., and Glöckner, F.O. 2012. The SILVA ribosomal RNA gene database project: improved data processing and web-based tools. Nucleic Acids Res 41(D1): D590–D596. doi:10.1093/nar/gks1219.

R Core Team. 2020. R: A Language and Environment for Statistical Computing. Vienna, Austria. Available from https://www.R-project.org/.

Rahal, S., and Chekireb, D. 2021. Diversity of rhizobia and non-rhizobia endophytes isolated from root nodules of Trifolium sp. growing in lead and zinc mine site Guelma, Algeria. Arch Microbiol 203(7): 3839–3849. doi:10.1007/s00203-021-02362-y.

Rahman, A., Manci, M., Nadon, C., Perez, I.A., Farsamin, W.F., Lampe, M.T., Le, T.H., Torres Martínez, L., Weisberg, A.J., Chang, J.H., and Sachs, J.L. 2023. Competitive interference among rhizobia reduces benefits to hosts. Current Biology 33(14): 2988–3001.e4. doi:10.1016/j.cub.2023.06.081.

Rajendran, G., Patel, M.H., and Joshi, S.J. 2012. Isolation and characterization of nodule-associated Exiguobacterium sp. from the root nodules of fenugreek (Trigonella foenum-graecum) and their possible role in plant growth promotion. Int J Microbiol. doi:10.1155/2012/693982.

Riley, A.B., Grillo, M.A., Epstein, B., Tiffin, P., and Heath, K.D. 2023. Discordant population structure among rhizobium divided genomes and their legume hosts. Mol Ecol 32(10): 2646–2659. John Wiley and Sons Inc. doi:10.1111/mec.16704.

Rosier, A., Beauregard, P.B., and Bais, H.P. 2021. Quorum Quenching Activity of the PGPR Bacillus subtilis UD1022 Alters Nodulation Efficiency of Sinorhizobium meliloti on Medicago truncatula. Front Microbiol 11. Frontiers Media S.A. doi:10.3389/fmicb.2020.596299.

Sprent, J. 1987. The ecology of the nitrogen cycle. Cambridge University Press.

Strauss, S.Y. 1991. Indirect effects in community ecology: Their definition, study and importance. Trends Ecol Evol 6(7): 206–210. doi:10.1016/0169-5347(91)90023-Q.

Strauss, S.Y., and Irwin, R.E. 2004. Ecological and Evolutionary Consequences of Multispecies Plant-Animal Interactions. Annu Rev Ecol Evol Syst 35(1): 435–466. doi:10.1146/annurev.ecolsys.35.112202.130215.

Tokgöz, S., Lakshman, D.K., Ghozlan, M.H., Pinar, H., Roberts, D.P., and Mitra, A. 2020. Soybean Nodule-Associated Non-Rhizobial Bacteria Inhibit Plant Pathogens and Induce Growth Promotion in Tomato. Plants 9(11): 1494. doi:10.3390/plants9111494.

Tsiknia, M., Tsikou, D., Papadopoulou, K.K., and Ehaliotis, C. 2020. Multi-species relationships in legume roots: From pairwise legume-symbiont interactions to the plant – microbiome - soil continuum. FEMS Microbiol Ecol. doi:10.1093/femsec/fiaa222.

Vitousek, P.M., Aber, J.D., Howarth, R.W., Likens, G.E., Matson, P.A., Schindler, D.W., Schlesinger, W.H., and Tilman, D.G. 1997. Human alteration of the global nitrogen cycle: sources and consequences. Ecological applications 7(3): 737–750.

Wagner, M.R., Lundberg, D.S., Coleman-Derr, D., Tringe, S.G., Dangl, J.L., and Mitchell-Olds, T. 2014. Natural soil microbes alter flowering phenology and the intensity of selection on flowering time in a wild Arabidopsis relative. Ecol Lett 17(6): 717–726. doi:10.1111/ele.12276.

Wagner, M.R., Lundberg, D.S., del Rio, T.G., Tringe, S.G., Dangl, J.L., and Mitchell-Olds, T. 2016. Host genotype and age shape the leaf and root microbiomes of a wild perennial plant. Nat Commun 7(1): 12151. doi:10.1038/ncomms12151.

Wendlandt, C.E., Regus, J.U., Gano-Cohen, K.A., Hollowell, A.C., Quides, K.W., Lyu, J.Y., Adinata, E.S., and Sachs, J.L. 2019. Host investment into symbiosis varies among genotypes of the legume Acmispon strigosus, but host sanctions are uniform. New Phytologist 221(1): 446–458. Blackwell Publishing Ltd. doi:10.1111/nph.15378.

Wickham, H. 2016. ggplot2: Elegant Graphics for Data Analysis. Springer International Publishing, New York, NY.

Wright, E.S. 2016. Using DECIPHER v2.0 to Analyze Big Biological Sequence Data in R. R J 8(1): 352. doi:10.32614/RJ-2016-025.

Yekutieli, D., and Benjamini, Y. 1999. Resampling-based false discovery rate controlling multiple test procedures for correlated test statistics. J Stat Plan Inference 82(1–2): 171–196. doi:10.1016/S0378-3758(99)00041-5.

Zilles, J.L., Rodríguez, L.F., Bartolerio, N.A., and Kent, A.D. 2016. Microbial community modeling using reliability theory. ISME J 10(8): 1809–1814. doi:10.1038/ismej.2016.1.

