## Supplementary text for "Co-inoculation with novel nodule-inhabiting bacteria reduces the benefits of legume-rhizobium symbiosis"

**SUPPLEMENTARY METHODS**

*Rhizobial and non-rhizobial strain isolation*

Strains were previously isolated from soils surrounding *M. truncatula* roots from 21 sites spanning the species’ native range: Spain, France, and Corsica. All nodules were surface sterilized prior to crushing and streaking for strain isolation. To determine the taxonomic identity of the isolated strains, Riley et al. (2023) performed several rounds of purification, extracted DNA from each purified culture, and submitted DNA for Illumina short-read shotgun sequencing. After trimming and quality filtering, Riley et al. (2023) conducted *de novo* assembly on the raw reads and submitted the resulting whole genome sequences to RASTtk for annotation (Brettin et al. 2015). While most strains were assigned to the expected *Sinorhizobium* spp., four were found to be “off-target” strains (522, 702A, 717A, and 733B), henceforth referred to as non-rhizobial endophytes (NREs). After determining that the four NREs were phylogenetically distant from rhizobia, based on colony morphology and a phylogeny constructed from several marker genes, Riley et al. (2023) excluded these strains from further analyses. All other data on these strains come from the present study.

*Taxonomic assignment with the Type (Strain) Genome Server*

The genome sequences for strains 522, 702A, 717A, and 733B were uploaded to the Type (Strain) Genome Server (TYGS) (Meier-Kolthoff and Göker 2019) for a whole genome-based taxonomic analysis. The analyses made use of recently introduced methodological updates and features (Meier-Kolthoff et al. 2022). Information on nomenclature, synonymy, and associated taxonomic literature was provided by TYGS’s sister database, the List of Prokaryotic names with Standing in Nomenclature (LPSN) (Meier-Kolthoff et al. 2022). Determination of closest type strain genomes was done in two complementary ways: First, all query NRE genomes were compared against all type strain genomes available in the TYGS database via the MASH algorithm (Ondov et al. 2016), and, the ten type strains with the smallest MASH distances chosen per user genome. Second, an additional set of ten closely related type strains was determined via the 16S rNA gene sequences. These were extracted from the user genomes using RNAmmer (Lagesen et al. 2007) and each sequence was subsequently BLASTed (Camacho et al. 2009) against the 16S rDNA gene sequence of each of the currently 20,220 type strains available in the TYGS database. This was used as a proxy to find the best 50 matching type strains (according to the bitscore) for each user genome and to subsequently calculate precise distances using the Genome BLAST Distance Phylogeny approach (GBDP) under the algorithm ‘coverage’ and distance formula *d_5_* (Meier-Kolthoff et al. 2013). These distances were finally used to determine the 10 closest type strain genomes for each of the user genomes.

*Greenhouse conditions and harvest*

We randomized four racks containing five plants per inoculation treatment (and control) (see Methods) across two greenhouse benches, resulting in 40 racks total (10 treatments x 4 replicate racks x 5 replicate plants = 200 plants), plus one additional rack of ten external control plants placed outside of the benches in the same room (210 total plants). To further reduce cross-contamination between treatments, immediately after inoculation, we added ½ cm of sterile sand to the surface of the soil of each Cone-tainer to provide a barrier between the inoculum and air. We programmed misters located inside the greenhouse room to go off for a duration of 45 minutes, four times a day.

Upon harvest (four weeks post-inoculation, see Methods), we cut the above-ground shoots of all plants at their bases and separately stored them in five-inch coin envelopes, which were left to dry in a 60 °C oven for three days before the dry-mass of each shoot was recorded. We extracted roots from Cone-tainers, dunked them into a bin of tap water to gently remove potting media, and then wrapped roots in paper towels and stored them in a 4 °C refrigerator until nodule dissection the following day. After counting the total number of nodules formed on each root, we haphazardly selected ten nodules, removing them carefully with forceps and weighing all ten together to the nearest 0.01 mg using a microbalance (Mettler-Toledo, Columbus OH, USA). After nodule dissection, we placed roots into five-inch coin envelopes, allowed them to dry using the same protocol for shoots, and recorded their weight.

*Nitrogen addition experiment growth conditions*

After scarification and surface-sterilization, we placed *Medicago* seeds on sterile petri dishes with sterile water and allowed them to germinate for 48 hours in the dark. Seedlings were then transplanted into magenta boxes (PlantMedia, Dublin, OH) containing autoclaved-sterilized 1:1 calcined clay and sand mixture to exclude any plant-available nitrogen. Plants were placed in a growth chamber at a constant 60% humidity with 16-hour, 23 °C days and 8-hour, 18 °C nights. One week after transplanting, plants were inoculated with each NRE individually or with sterile media. Plants received 5 mL of N-supplemented Fahräeus media (Vincent, 1970) once a week for four weeks after inoculation. We used ammonium nitrate (NH_4_NO_3_) as the external N source, ramping up the concentration at each application to reflect varying nitrogen requirements during seedling development (Barker et al., 2006): 1.0 mg/mL, 2.0 mg/mL, 3.0 mg/mL, and 5.0 mg/mL, week-by-week for four weeks until harvest.

*Tissue occupancy experiment conditions*

We grew ten plants of each inoculation treatment for tissue occupancy (see Methods) in pre-sterilized plastic, drainless, three-inch plant trays that were covered with transparent, plastic 10-inch domes, yielding 50 plants in total. We filled plant trays with sterile tap water, so that plants would receive a consistent supply of water. We used the same inoculation methods as those described for the nitrogen addition experiment, but after rinsing roots, we placed them in a sterile 50 mL falcon tube with 40 mL sterile DI water and shook thoroughly to remove excess soil and weakly-associated rhizosphere microbes. After washing roots this way, we placed each root into its own sterile 50 mL falcon tube with silica beads and cotton at the bottom (to remove excess moisture) and left to dry overnight. We removed nodules from roots as well as root tissue sections without nodules. Nodules and roots were placed in separate 1.5 mL tubes with 500 μL sterile water and allowed to re-imbibe overnight.

**SUPPLEMENTARY RESULTS**

*Taxonomy*

The resulting *Paenibacillus* tree inferred by TYGS (Figure 1A) placed *Paenibacillus* sp. 522 in a species cluster with no other taxa, as a sister taxon to *Paenibacillus silvestris* 5J-6 (dDDH *d_4_* = 48.1%) with 100 percent bootstrap support at this node. The *Bacillus* tree (Figure 1B) placed *Bacillus* sp. 717A in its own species cluster and as a sister taxon to *Bacillus safensis* FO-36b (dDDH *d_4_* = 68.4%) with 59 percent bootstrap support. The *Pseudomonas* tree (Figure 1C) placed *Pseudomonas* sp. 702A at the node sister eight *Pseudomonas* type strains with 100 percent bootstrap support, of which the closest neighbor was *Pseudomonas hormoni* G20-18T (dDDH *d_4_* = 33.5%). *Pseudomonas* sp. 733B was placed with 100 percent bootstrap support sister to a clade with two *Pseudomonas* type strains, of which the closest neighbor was *Pseudomonas azerbaijanoccidentalis SWRI74* (dDDH *d_4_* = 39.1%). Both *Pseudomonas* sp. 702A and *Pseudomonas* sp. 733B were placed in their own, separate, species clusters with no other taxa. For a list of strain information present in the trees in Figure 1 as well as their calculated genome distances, see Tables S1-S2.

*Identification of NRE strains among amplicon sequence variants*

After aligning our inferred ASVs against the 16S rRNA sequences of our isolate genomes, we found that ASV1 and ASV5 aligned perfectly to the *Sinorhizobium* and *Pseudomonas* sp. 733B 16S V3-V4 regions, respectively, retaining 100% of the ASV lengths in the alignments (Table S12). Our taxonomic assignments of ASVs based on alignments to SILVA are consistent with these predictions (Figure 4). No ASVs aligned perfectly to the *Bacillus* sp. 717A 16S gene (Table S9). Among the ASVs that did align to *Bacillus* sp. 717A, the most abundant ASV in our dataset was ASV28 (Table S9; Table S7; Figure 4), which aligned at a 94% identity with 100% coverage of the ASV (Table S9). Several *Bacillus* species have displayed relatively high levels of intraspecific sequence heterogeneity in their 16S rRNA genes (Coenye and Vandamme 2003), which likely explains this smaller alignment identity compared to our other inoculum strains. This sequence was indeed assigned to the genus *Bacillus* based on the SILVA database (Figure 4), and, overall, out of all other ASVs inferred from the amplicon data, ASV28 is the most likely representative of *Bacillus* sp. 717A.

**REFERENCES**

Benjamini, Y., and Hochberg, Y. 1995. Controlling the False Discovery Rate: A Practical and Powerful Approach to Multiple Testing. Journal of the Royal Statistical Society: Series B (Methodological) **57**(1): 289–300. doi:10.1111/j.2517-6161.1995.tb02031.x.

Brettin, T., Davis, J.J., Disz, T., Edwards, R.A., Gerdes, S., Olsen, G.J., Olson, R., Overbeek, R., Parrello, B., Pusch, G.D., Shukla, M., Thomason, J.A., Stevens, R., Vonstein, V., Wattam, A.R., and Xia, F. 2015. RASTtk: A modular and extensible implementation of the RAST algorithm for building custom annotation pipelines and annotating batches of genomes. Sci Rep **5**. doi:10.1038/srep08365.

Brown, S., Grillo, M., Podowski, J., and Heath, K.D. 2020. Soil origin and plant genotype structure distinct microbiome compartments in the model legume Medicago truncatula. Microbiome **8**(139). doi:10.21203/rs.3.rs-21175/v1.

Callahan, B.J., McMurdie, P.J., Rosen, M.J., Han, A.W., Johnson, A.J.A., and Holmes, S.P. 2016. DADA2: High-resolution sample inference from Illumina amplicon data. Nat Methods **13**(7): 581–583. doi:10.1038/nmeth.3869.

Camacho, C., Coulouris, G., Avagyan, V., Ma, N., Papadopoulos, J., Bealer, K., and Madden, T.L. 2009. BLAST+: architecture and applications. BMC Bioinformatics **10**(1): 421. doi:10.1186/1471-2105-10-421.

Coenye, T., and Vandamme, P. 2003. Intragenomic heterogeneity between multiple 16S ribosomal RNA operons in sequenced bacterial genomes. FEMS Microbiol Lett **228**(1): 45–49. doi:10.1016/S0378-1097(03)00717-1.

Lagesen, K., Hallin, P., Rødland, E.A., Stærfeldt, H.-H., Rognes, T., and Ussery, D.W. 2007. RNAmmer: consistent and rapid annotation of ribosomal RNA genes. Nucleic Acids Res **35**(9): 3100–3108. doi:10.1093/nar/gkm160.

Love, M.I., Huber, W., and Anders, S. 2014. Moderated estimation of fold change and dispersion for RNA-seq data with DESeq2. Genome Biol **15**(12): 550. doi:10.1186/s13059-014-0550-8.

Meier-Kolthoff, J.P., Auch, A.F., Klenk, H.-P., and Göker, M. 2013. Genome sequence-based species delimitation with confidence intervals and improved distance functions. BMC Bioinformatics **14**(1): 60. doi:10.1186/1471-2105-14-60.

Meier-Kolthoff, J.P., Carbasse, J.S., Peinado-Olarte, R.L., and Göker, M. 2022. TYGS and LPSN: a database tandem for fast and reliable genome-based classification and nomenclature of prokaryotes. Nucleic Acids Res **50**(D1): D801–D807. doi:10.1093/nar/gkab902.

Meier-Kolthoff, J.P., and Göker, M. 2019. TYGS is an automated high-throughput platform for state-of-the-art genome-based taxonomy. Nat Commun **10**(1): 2182. doi:10.1038/s41467-019-10210-3.

Ondov, B.D., Treangen, T.J., Melsted, P., Mallonee, A.B., Bergman, N.H., Koren, S., and Phillippy, A.M. 2016. Mash: Fast genome and metagenome distance estimation using MinHash. Genome Biol **17**(1). doi:10.1186/s13059-016-0997-x.

Riley, A.B., Grillo, M.A., Epstein, B., Tiffin, P., and Heath, K.D. 2023. Discordant population structure among rhizobium divided genomes and their legume hosts. Mol Ecol **32**(10): 2646–2659. John Wiley and Sons Inc. doi:10.1111/mec.16704.

Yekutieli, D., and Benjamini, Y. 1999. Resampling-based false discovery rate controlling multiple test procedures for correlated test statistics. J Stat Plan Inference **82**(1–2): 171–196. doi:10.1016/S0378-3758(99)00041-5.

**SUPPLEMENTARY FIGURES**

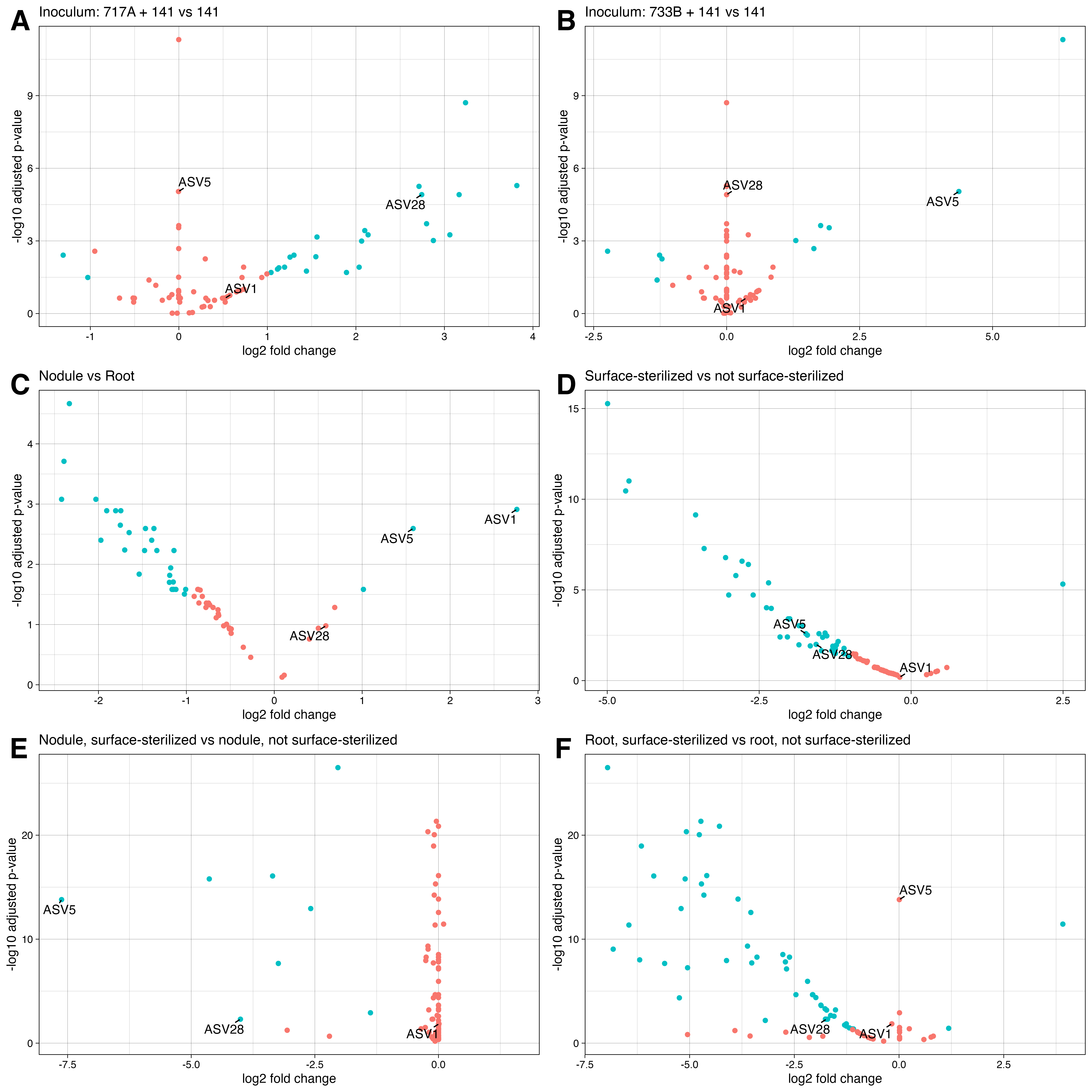

**Figure S1. Differential abundance of ASVs across inocula, tissue sections, and surface-sterilization treatments.** Using a minimum fold change 1.5 and *P* (false-discovery rate adjusted) < 0.05, amplicon sequence variants (ASVs) diferentially abundant across contrasts of inocula, tissues, and surface-sterilization treatments were inferred. Points refer to individual ASVs, blue points correspond to signifiicant ASVs while red points are insignificant. The second term in plot titles indicate baselines in comparisons. ASVs representing inocula strains are labeled, ASV1 = *S. meliloti* 141; ASV28 = *Bacillus* sp. 717A; ASV5 = *Pseudomonas* sp. 733B. (A) ASV28 was enriched in *Bacillus* sp. 717A + *S. meliloti* 141 co-inoculated samples compared to samples inoculated with *S. meliloti* 141 only. Similarly, (B) ASV5 was enriched in *Pseudomonas* sp. 733B + *S. meliloti* 141 co-inoculated samples compared to samples inoculated with *S. meliloti* 141 only. (C) Both ASV5 and ASV1 were enriched in nodule samples compared to root samples. (D) ASV5 and ASV28 were enriched in non-surface-sterilized samples compared to surface-sterilized samples, while ASV1 was not differentially abundant across either group. (E) Among just nodule samples, ASV5 and ASV28 were enriched in non-surface-sterilized samples while ASV1 was not differentially abundant. (F) Among just root samples, only ASV28 was enriched in non-surface-sterilized root samples while ASV1 and ASV5 were not different across either group.
